# HURP facilitates spindle assembly by stabilizing microtubules and working synergistically with TPX2

**DOI:** 10.1101/2023.12.18.571906

**Authors:** Venecia Valdez, Meisheng Ma, Bernardo Gouveia, Rui Zhang, Sabine Petry

## Abstract

In large vertebrate spindles, the majority of microtubules are formed via branching microtubule nucleation, whereby microtubules nucleate along the side of pre-existing microtubules. Hepatoma up-regulated protein (HURP) is a microtubule-associated protein that has been implicated in spindle assembly, but its mode of action is yet to be defined. In this study, we show that HURP is necessary for RanGTP-induced branching microtubule nucleation in *Xenopus* egg extract. Specifically, HURP stabilizes the microtubule lattice to promote microtubule formation from γ- TuRC. This function is shifted to promote branching microtubule nucleation in the presence of TPX2, another branching-promoting factor, as HURP’s localization to microtubules is enhanced by TPX2 condensation. Lastly, we provide a structure of HURP on the microtubule lattice, revealing how HURP binding stabilizes the microtubule lattice. We propose a model in which HURP stabilizes microtubules during their formation, and TPX2 preferentially enriches HURP to microtubules to promote branching microtubule nucleation and thus spindle assembly.

## Introduction

Cell division relies on the formation of a bipolar spindle to orient and faithfully segregate chromosomes into two daughter cells. The spindle is made up of many microtubules (MTs), filaments composed of tubulin dimers, that must be nucleated and organized in a regulated manner. Additionally, MT dynamics (e.g., growth rate, catastrophe, and rescue) are regulated by MT-associated proteins (MAPs) to facilitate proper spindle assembly. Dysregulation of MT nucleation, dynamics, and organization can lead to improper chromosome segregation and result in apoptosis or diseases such as cancer^1^. Therefore, dissecting the complexity of how MTs are formed and regulated is crucial to understanding how cell division is continuously executed in a reliable manner.

In large vertebrate spindles, branching MT nucleation forms most spindle MTs^2–5^. In this process, new MTs nucleate at a shallow angle on the side of pre-existing MTs, resulting in exponential self-amplification^6^. In vitro, the minimal branching components are tubulin, the gamma-tubulin ring complex (γ-TuRC) which serves as the universal microtubule nucleation template, and the augmin complex which localizes γ-TuRC to a pre-existing microtubule^7–9^. In *Xenopus laevis* egg extract, the general nucleation factor XMAP215/chTOG and the protein TPX2 is also required for branching microtubule nucleation^6,10^. TPX2 forms a co-condensate with tubulin on a MT and recruits the augmin complex, which in turn recruits γ-TuRC^7,11,12^. Branching MT nucleation is regulated via the small GTPase Ran. Both the augmin complex and TPX2 are spindle assembly factors (SAF), meaning that their binding to MTs is inhibited by binding of importins and released by RanGTP^13,14^. Given that RanGTP is present in a gradient centered around chromatin, branching MT nucleation is spatially regulated to promote nucleation near chromosomes^5,15,16^.

Here, we hypothesized that the hepatoma upregulated protein (HURP) plays a role in branching MT nucleation. HURP is another SAF^17,18^, first identified in human hepatocellular carcinoma, and has been implicated in a variety of cancers as a putative oncogene^19–23^. Studies in human cells have demonstrated that HURP is a MT-bundling protein that localizes predominantly to kinetochore fibers (k-fibers) and is important in k-fiber stability and chromosome congression^18,24,25^. Interestingly, in acentrosomal systems, such as *Drosophila* oocytes, mouse oocytes, and *Xenopus* egg extract, loss of HURP leads to spindle assembly defects and a reduction of spindle MT density^17,26–28^. Additionally, HURP directly or indirectly interacts with TPX2 and augmin subunits in various species^28–30^, which prompted us to investigate HURP’s role in branching microtubule nucleation.

In this work, we uncover that HURP is integral to branching MT nucleation. By combining live imaging in Xenopus egg extract with in vitro assays and single particle cryo-electron microscopy (cryo-EM), we reveal how HURP promotes microtubule formation. Altogether, this work uncovers how HURP promotes branching microtubule nucleation and thereby spindle assembly.

## Results

### HURP is required for branching MT nucleation in *Xenopus* egg extract

To investigate whether HURP plays a role in branching MT nucleation, we first developed a strategy to remove HURP from *Xenopus* egg extract. We generated a HURP-specific antibody and coupled it to beads. The conjugated beads were incubated with egg extract and then removed to immunodeplete endogenous HURP (present at ∼320 nM^31^) (**Fig. 1a**). When compared to mock-depleted extract, Western blot analysis revealed no co-depletion of other MT nucleation and branching factors such as γ-tubulin, XMAP215, TPX2, and augmin in the HURP-depleted extract. This indicates that the immunodepletion was specific and HURP does not strongly associate with any of these proteins under these conditions. This is consistent with previous work suggesting that HURP interacts with TPX2 and XMAP215 in a RanGTP-dependent manner^28^.

**Figure 1:**
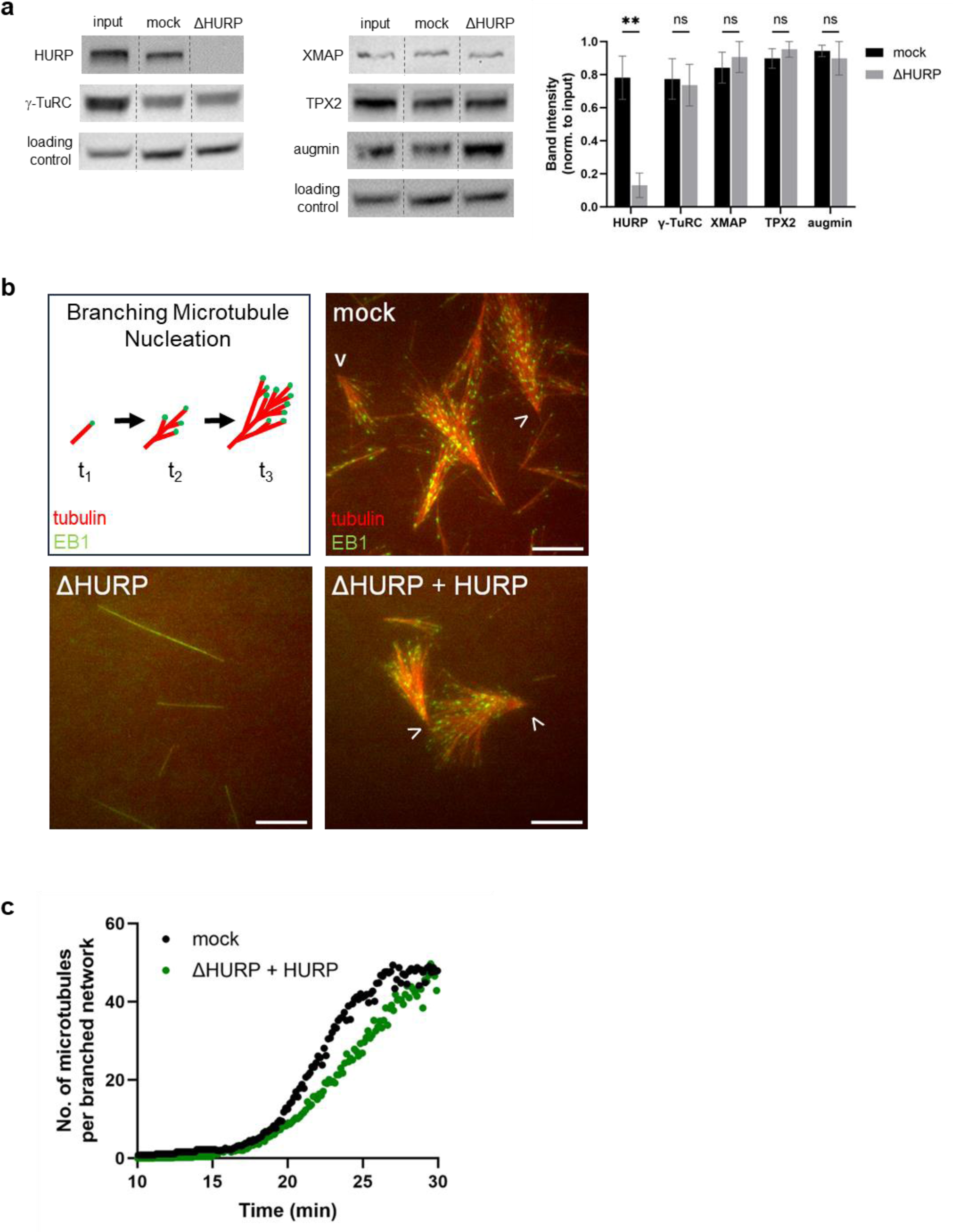
HURP is necessary for branching microtubule nucleation in Xenopus egg extract. **a**Western blot of *Xenopus* egg extract input, mock-depleted extract, and HURP-depleted extract probed for HURP, γ-TuRC (γ-tubulin), TPX2, and augmin (HAUS1). α-tubulin was probed as a loading control. Rearrangement of nonadjacent lanes (for display purposes) are indicated by dotted lines. Band intensities, normalized to the input lane, were averaged across four replicates and plotted in the bar graph (mean±SEM) for mock-depleted and HURP-depleted extract. Significance is defined as a p-value < 0.5 determined by two-tailed unpaired Student’s t-test. ns = not significant. Double asterisks (**) indicate a p-value < 0.01 **b** RanGTP-induced branching microtubule (MT) nucleation assay in mock-depleted extract, HURP-depleted extract, and HURP-depleted extract plus addback of 250nM purified HURP construct. MTs (AlexaFluor647-tubulin, pseudo-colored red) and mCherry-EB1 (pseudo-colored green) were imaged for each reaction over 30 minutes. Representative cropped images (51.54 μm x 51.54 μm) at a timepoint of ∼22 minutes are displayed. Scale bar = 10 µm **c** Plot of MT number (EB1 foci) over time (min) in a 110.03 μm x 93.41 μm crop from the movies represented in panel b, normalized by the number of branched structures present at the ∼25 minute timepoint. The mock depletion contained 30 fans. The HURP-depleted plus addback of 250nM purified HURP contained 9 fans. Branching MT nucleation assays were repeated three times with three separate egg extracts and all portrayed similar results. A representative example from one replicate is displayed in this figure.

Next, we directly observed branching MT nucleation in the presence and absence of HURP in *Xenopus* egg extract using total internal reflection fluorescence (TIRF) microscopy. In this assay, MTs are visualized via fluorescent tubulin and their growing plus ends via fluorescent end-binding protein 1 (EB1)^6^. Branching MT nucleation is initiated by the addition of constitutively active RanGTP(Q69L), while addition of the ATPase inhibitor vanadate inhibits motor activity to prevent reorganization of the resulting fan-like branched MT networks. As expected, the mock-depleted extract yielded branched, fan-like MT structures in the presence of RanGTP(Q69L). To our surprise, no branched MT structures formed in the absence of HURP (**Fig. 1b and Supplementary Movie 1**). Instead, a few single MTs were nucleated and polymerized, but branched MTs no longer formed. To verify that the observed phenotype was specific for HURP, we purified full-length HURP and added it back to HURP-depleted extract, which rescued the formation of branched MT structures and thus indicates that the observed phenotype is a direct consequence of HURP depletion. Although the addition of 250 nM of purified HURP rescued the formation of branched MT structures, the rescue yielded fewer total fans than the mock depleted extract (**Supplementary Fig. S1)**. This may be due to adding back sub-endogenous levels of HURP (250 nM) or an inactive fraction in the final purified HURP eluate. We therefore needed to determine, of the individual branched networks that formed, if the MT nucleation kinetics were similar between the mock and rescue conditions. We measured the number of MTs (counted as EB1 foci) over time normalized to the number of fans present (**Supplementary Fig. S2**) and found that the rescue assay had similar nucleation kinetics as the mock depletion (**Fig. 1c**). In conclusion, HURP is necessary for branching MT nucleation in *Xenopus* egg extract.

### HURP promotes MT formation and works together with TPX2

Some MAPs, such as XMAP215 and TPX2, are required for and induce single MT nucleation or branching MT nucleation, respectively, when added in excess to meiotic *Xenopus* egg extract^6,10^. To assess the effect of HURP on MT formation, we added increasing amounts of purified HURP to *Xenopus* egg extract and visualized the reaction via TIRF microscopy. We find that addition of purified HURP (125 nM and 250 nM excess) promotes the formation of single MTs in a dose-dependent manner (**Figs. 2a, b, Supplementary Movie 2, and Supplementary Fig. S3)**. The formation of single MTs is similar to what is observed in the presence of increasing XMAP215 concentration^10^, but differs from the formation of branched MT networks observed when an excess of TPX2 is added to extract^6,32^. The different outcomes of excess HURP and excess TPX2 are interesting given that both HURP and TPX2 are SAFs^18,30^. This discrepancy suggests that HURP is working in a manner distinct from TPX2 to form MTs.

**Figure 2:**
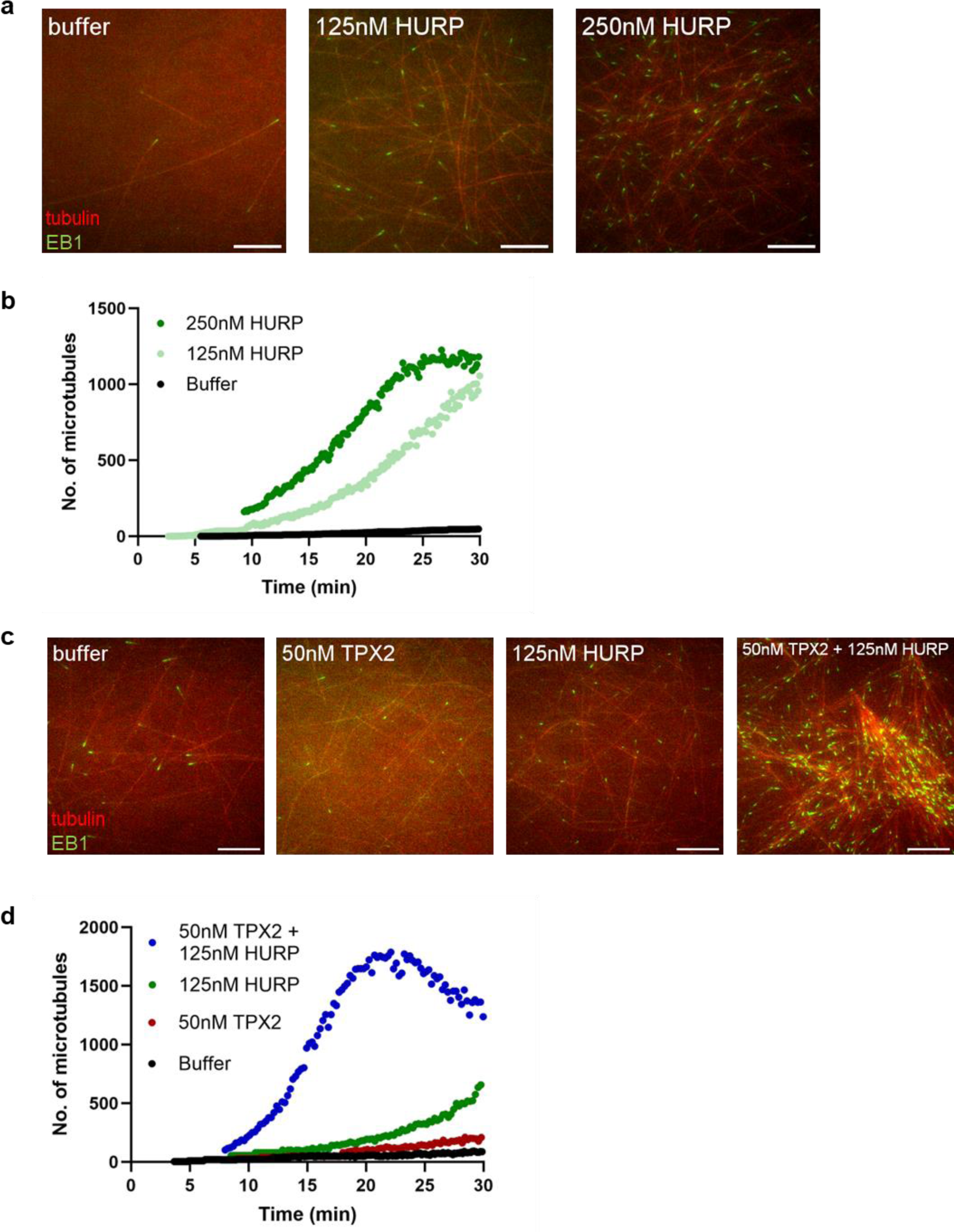
Excess HURP induces microtubule nucleation in Xenopus egg extract. **a**Microtubule (MT) nucleation assay in *Xenopus* egg extract induced by addition of purified HURP at various concentrations (no added RanGTP). Addition of CSF-XB buffer served as the negative control. MTs (AlexaFluor647-tubulin, pseudo-colored red) and mCherry-EB1 (pseudo-colored green) were imaged for each reaction over 30 minutes. Representative cropped images (51.54 μm x 51.54 μm) at a timepoint of ∼15 minutes are displayed. Scale bar = 10 µm **b** Plot of MT number (EB1 foci) over time (min) in a 110.03 μm x 93.41 μm crop from the movies represented in panel a. **c** MT nucleation assay in Xenopus egg extract induced by adding purified 50 nM TPX2, 125 nM HURP, or both (no added RanGTP). Addition of CSF-XB buffer served as the negative control. MTs and EB1 were imaged for each reaction over 30 minutes. Representative cropped images (51.54 μm x 51.54μm) at a timepoint of ∼15 minutes are displayed. The brightness and contrast for the buffer condition was adjusted individually for display purposes. Scale bar = 10 µm **d** Plot of MT number (EB1 foci) over time (min) in a 110.03 μm x 93.41 μm crop from the movies represented in panel c. All MT nucleation assays were repeated three times with three separate egg extracts and all portrayed similar results. A representative example from one replicate for each experiment is displayed in this figure.

To further understand the relationship between HURP and TPX2, we assessed whether HURP and TPX2 work together to promote the formation of MTs. A low level of excess TPX2 alone (50 nM excess) is not sufficient to promote the formation of branched MT networks, whereas 125 nM of HURP induces the formation of single MTs. In contrast, when we added HURP to *Xenopus* egg extract containing a low level of excess TPX2, we observed a large increase in the number of MTs formed (**Figs. 2c, 2d, Supplementary Movie 3, and Supplementary Fig. S4**). In fact, the number of MTs formed over time increased exponentially, a hallmark of branching MT nucleation^6^, and some branched MT networks could be observed. This suggests that the presence of TPX2 influences the mechanism through which HURP promotes MT formation. In addition, this data suggests that HURP enhances the ability of TPX2 to promote the formation of branched MT networks.

### TPX2 recruits additional HURP to MTs in vitro

To investigate how TPX2 and HURP influence each other on the MT, we performed sequential MT binding reactions in vitro. We find that when GMPCPP-stabilized MTs are pre-incubated with low levels of TPX2 (50 nM), the subsequent binding of HURP to the MT is unaffected when compared to MTs pre-incubated with BRB80 buffer (**Figs. 3a and 3b**). Similarly, pre-incubation of MTs with HURP does not affect the subsequent recruitment of TPX2 to the MT (**Figs. 3c and 3d**).

**Figure 3:**
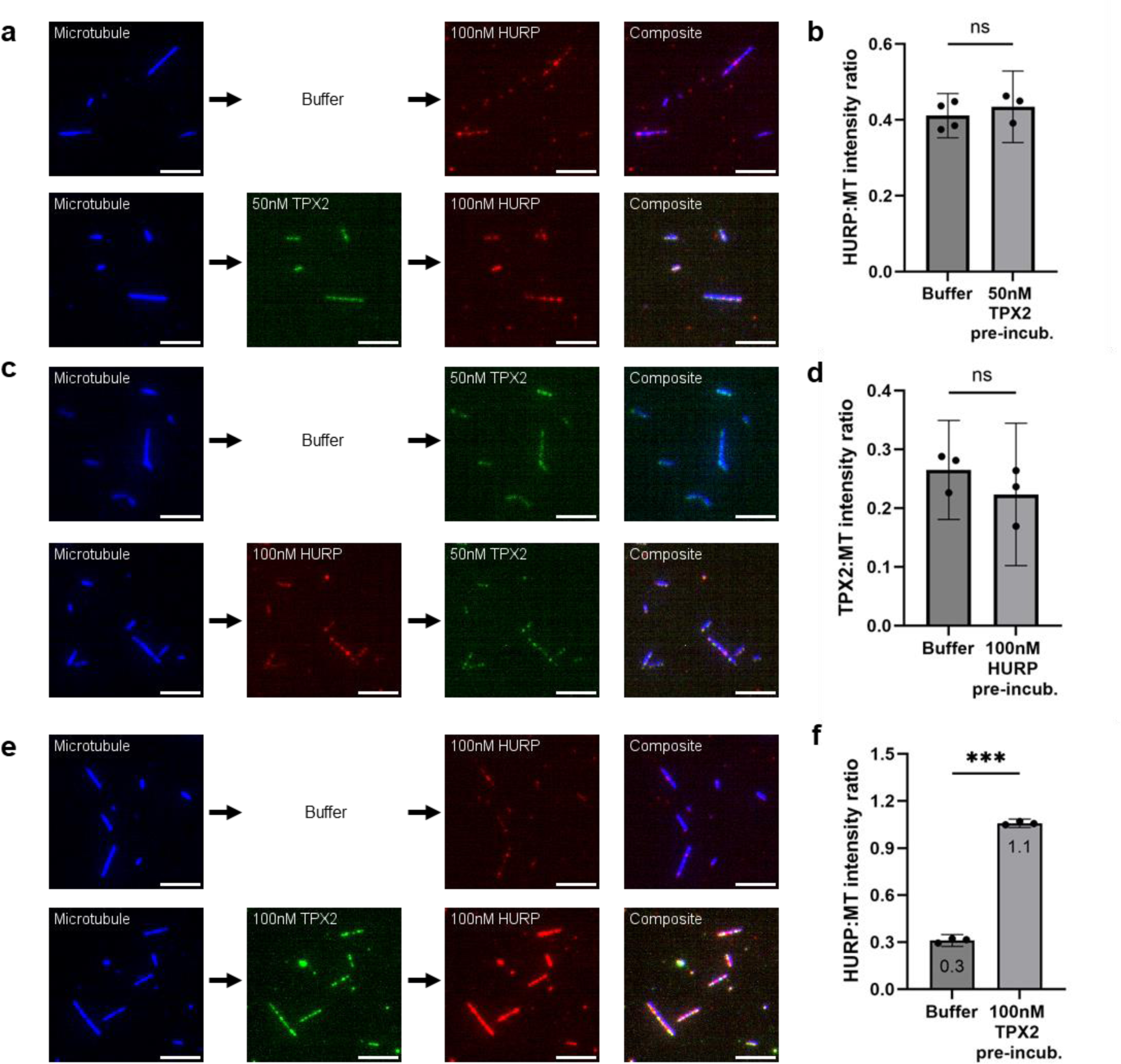
TPX2 enriches HURP to microtubules in vitro. Sequential in vitro microtubule (MT) binding reactions using biotinylated, GMPCPP-stabilized MTs (ATTO647N-tubulin, pseudocolored blue) bound to functionalized coverslips. Arrows between images in panels a, c, and e indicate the sequence of the MT binding reaction for each experimental condition. Each row of images portrays one experimental condition. Representative cropped images (39.77 µm x 39.77 µm) are displayed. Scale bars = 5 µm **a** MTs were pre-incubated with 50 nM GFP-TPX2 (n=3) or CSF-XB buffer (negative control, n=4), washed, then bound with 100 nM mCherry-HURP. MTs and bound proteins were imaged after a final BRB80 wash. **b** Bar graph plotting the average HURP:MT intensity ratio across replicates for each sequential MT binding reaction in panel a. Individual points represent a single replicate (>25 MTs per replicate). Error bars represent the 95% confidence interval **c** MTs were pre-incubated with 100 nM mCherry-HURP (n=3) or CSF-XB buffer (negative control, n=3), washed, then bound with 50 nM GFP-TPX2. MTs and bound proteins were imaged after a final BRB80 wash. **d** Bar graph plotting the average TPX2:MT intensity ratio across replicates for each reaction in panel c. Individual points represent a single replicate (>25 MTs per replicate). Error bars represent the 95% confidence interval **e** MTs were pre-incubated with 100 nM GFP-TPX2 (n=3) or CSF-XB buffer (negative control, n=3), washed, then bound with 100 nM mCherry-HURP. MTs and bound proteins were imaged after a final BRB80 wash. **f** Bar graph plotting the average HURP:MT intensity ratio across replicates for each reaction in panel e. Averages are denoted within the bar for each condition. Individual points represent a single replicate (>25 MTs per replicate). Error bars represent the 95% confidence interval. Significance for all bar plots is defined as p-value < 0.5 and determined by Welch’s two-tailed t-test. ns = not significant. Triple asterisks (***) represent a p-value < 0.001

Because TPX2 enhances the formation of branched MT structures by forming a biomolecular condensate on the MT surface, which recruits soluble tubulin^11^, we hypothesized that condensed TPX2 is capable of enhancing the localization of HURP to the MT. The phase boundary above which TPX2 forms condensates on MTs is ∼100 nM^33^. When MTs were pre-incubated with 100 nM TPX2, we observed that subsequent localization of HURP increased by more than three-fold (**Figs 3e and 3f**). This data suggests that condensed TPX2 can specifically enrich HURP on the MT.

To test whether this relationship also exists in solution, we performed an in vitro bulk phase separation assay, in which TPX2 forms condensates in the absence of MTs. Unlike TPX2, HURP at high concentrations of up to 1 µM in low-salt buffer (BRB80) is incapable of forming condensates (Supplementary Fig. 5a). Furthermore, in the presence of equimolar tubulin, which lowers the phase boundary of TPX2 to 50 nM^11^, HURP remains incapable of forming condensates (Supplementary Fig. 5b). However, HURP was recruited to TPX2 condensates in solution (**Supplementary Fig. 5b**). Altogether, these data show that HURP is a client of TPX2 condensates. Because TPX2 condensates preferentially form on MTs, this in turn enhances localization of HURP to the MT.

### HURP facilitates MT formation from γ-TuRC in vitro by stabilizing the MT lattice

Having established that HURP is required for Ran-induced branching MT nucleation and that it promotes the formation of single MTs in *Xenopus* extract, we next addressed how HURP accomplishes these actions. Because nucleation of single MTs requires γ-TuRC as the universal MT template^34^, we directly investigated how HURP influences MT nucleation from γ-TuRC. In our minimal in vitro assay, biotinylated γ-TuRC is tethered to the surface of a biotinylated and passivated glass coverslip via neutravidin. Upon introducing fluorescently labeled tubulin, the formation of MTs from γ-TuRC can be observed in real time using TIRF microscopy^35–37^. Interestingly, when 50 nM HURP was added, we observed a ∼3-fold increase in the number of γ- TuRC-mediated MTs nucleated in comparison to the buffer control (**Figs 4a, b, and Supplementary Movie 4**). By averaging the number of MTs over time, and fitting to an exponential function, we obtained nucleation rates and observed a ∼2.4-fold increase in the presence of 50 nM HURP (0.47 MTs/sec) in comparison to the buffer control (0.20 MTs/sec). Thus, HURP directly increases the number of MTs nucleated from γ-TuRC.

**Figure 4:**
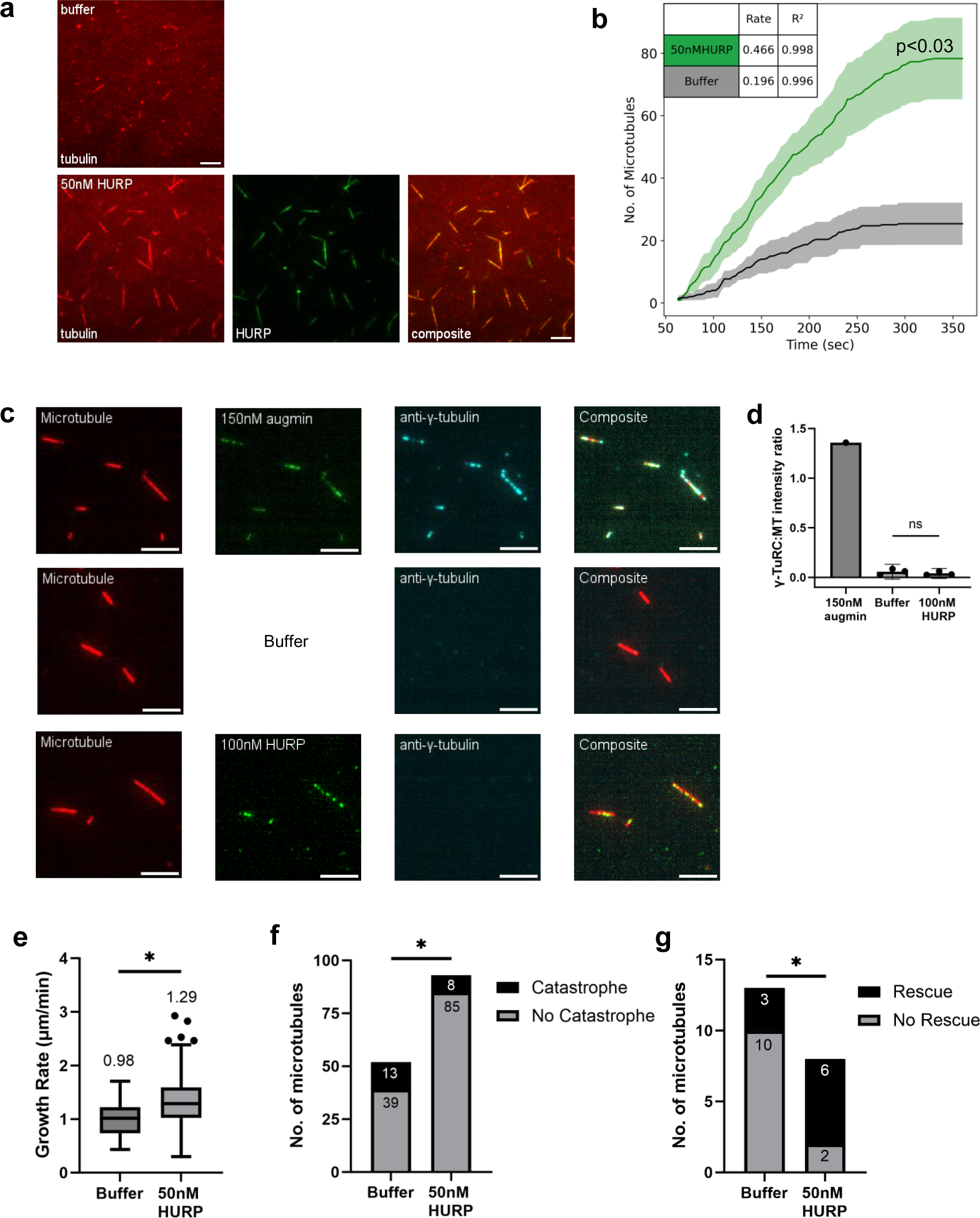
HURP facilitates nucleation from γ-TuRC in vitro and stabilizes the microtubule lattice. **a**In vitro γ-TuRC microtubule (MT) nucleation assay in the presence of 10 μM Alexa Fluor 568- tubulin plus BRB80 buffer (negative control, n=3) or 50 nM GFP-HURP (n=4). Still images at the final timepoint (6 min). Representative cropped images (39.77 µm x 39.77 µm) displayed. Scale bar = 5 µm. **b** Line graph depicting average MT number over time (sec) ± SEM for each condition in panel a. Nucleation rates (MTs per sec, denoted in inset) calculated by fitting data to an exponential function. P-value calculated by comparing the final timepoint of the buffer (25±6.7 MTs; mean±SEM) and 50 nM HURP (78±13 MTs) conditions via two-tailed Student’s t-test. **c** In vitro microtubule (MT) binding reactions using biotinylated, GMPCPP-stabilized MTs (Alexa Fluor 568-tubulin) bound to functionalized coverslips. Each row of images portrays one experimental condition. MTs were incubated with a mix of 70 nM γ-TuRC and either 150 nM GFP-augmin (positive control, n=1), CSF-XB buffer (negative control, n=3), or 100 nM GFP-HURP (n=3) in CSFXB buffer. Alexa Fluor 647-conjugated γ-tubulin antibody was used to visualize γ-TuRC (pseudo-colored cyan). MTs and bound proteins were imaged after a final BRB80 wash. Representative cropped images (19.33 µm x 19.33 µm) displayed. Scale bar = 5 µm **d** Bar graph plotting average γ-TuRC(γ-tubulin):MT intensity ratio for panel d. Individual points represent replicates (>25 MTs per replicate). Significance calculated using Welch’s two-tailed t-test. **e** Box and whisker plot (Tukey) for MT growth rates (μm/min) from all microtubules across replicates from the in vitro γ-TuRC nucleation assay in panel a. Significance calculated by Welch’s two-tailed t-test using averages from each replicate per condition **f** Stacked bar graphs plotting the number of MTs undergoing catastrophe or not undergoing catastrophe across replicates for panel a. Significance between conditions was calculated by Fisher’s exact test. **g** Stacked bar graphs plotting the number of MTs being rescued or not rescued after a catastrophe event (recorded in panel f). Significance between conditions was calculated by Fisher’s exact test. Significance defined as a p-value < 0.05. ns = not significant. A single asterisk (*) indicates a p-value < 0.05.

To test if HURP acts directly on γ-TuRC, we performed a MT binding assay in vitro using GMPCPP-stabilized MTs. If HURP binds to γ-TuRC, we would expect higher γ-TuRC localization to MTs when γ-TuRC is pre-incubated with HURP than a buffer control. As a positive control, we performed this assay with augmin, which is known to directly bind both MTs and γ-TuRC^12^. As expected, we detected that pre-incubation of γ-TuRC with augmin results in localization of γ-TuRC to the MT, visualized by an AlexaFluor649-conjugated γ-tubulin antibody (**Figs. 4c and d**). Conversely, pre-incubation of γ-TuRC with HURP could not localize γ-TuRC to the MT. These results suggest that HURP does not directly bind to γ-TuRC to enhance MT nucleation.

We next asked whether HURP influences MT dynamics. Using the minimal in vitro assay with tethered γ-TuRC, we measured the MT growth rate. Indeed, the MT growth rate is slightly faster in the presence of HURP (1.29 ± 0.08 µm/min) when compared to the buffer control (0.98 ± 0.09 µm/min) (**Fig. 4e**). Because HURP has not previously been characterized as having a direct effect on polymerization rate, we investigated this result further. Canonical polymerases such as XMAP215 increase MT growth speed by localizing soluble tubulin to the MT plus-end via conserved TOG domains^38^. Here, we found that HURP is incapable of localizing soluble tubulin to a GMPCPP-stabilized MT lattice (**Supplementary Fig. S6**). This result, as well as the lack of TOG domains in the HURP sequence, suggests that HURP is functioning in a manner different from polymerases like XMAP215.

Strikingly, we detected a ∼3-fold reduction in the proportion of MTs experiencing catastrophes in the presence of HURP (8/93 MTs, ∼8.6%) compared to the buffer condition (13/52 MTs, ∼25%) (**Fig. 4f**). Additionally, the subsequent proportion of MTs rescued was also significantly increased (∼3-fold) in the HURP condition (6/8 catastrophes rescued, 75%) in comparison to the buffer (3/13 catastrophes rescued, 23%) (**Fig. 4g**). These results show that HURP confers stability to MTs. This is consistent with previous studies characterizing cold-resistance of k-fibers in HURP-depleted cells, in that loss of HURP resulted in less stable (cold-resistant) k-fibers^18^. Altogether, HURP stabilization of nascent MTs may result in the increase of optically observable MTs over time in our in vitro assay with γ-TURC nucleated MTs.

### A bipartite microtubule binding mode of HURP

To better understand how HURP binding to the MT lattice influences MT dynamics, we performed single particle cryo-EM of HURP bound to GMPCPP-stabilized MT. Following a previous study^39^, we made a human HURP^65-174^ construct that covers the MTBD1 and a HURP^1-300^ construct that covers both MTBD1 and MTBD2, two regions that are highly conserved between human and Xenopus laevis (**Figs. 5a, 5b, and Supplementary Fig. S7**). We added these two protein constructs, HURP^65-174^ and HURP^1-300^, to GMPCPP-stabilized MTs in an over-saturated condition, and determined their cryo-EM structures using a newly developed MT data processing pipeline based on CryoSPARC^40^ (**Supplementary Fig. S8**) that utilized the results from MT seam search^41^. Both structures (HURP^65-174^ and HURP^1-300^) revealed nearly identical MAP densities (**Figs. 5c and 5d**). The structure of the HURP^65-174^-decorated MT at 2.8 Å local resolution allowed us to perform unbiased and automated protein identification of the non-tubulin densities using the software ModelAngelo^42^. Impressively, a search of the ModelAngelo-derived protein sequence against the entire human proteome produced a single hit of HURP protein with high confidence. The hit was further verified by careful visual inspection of the fit between the atomic model and cryo-EM density map.

**Figure 5:**
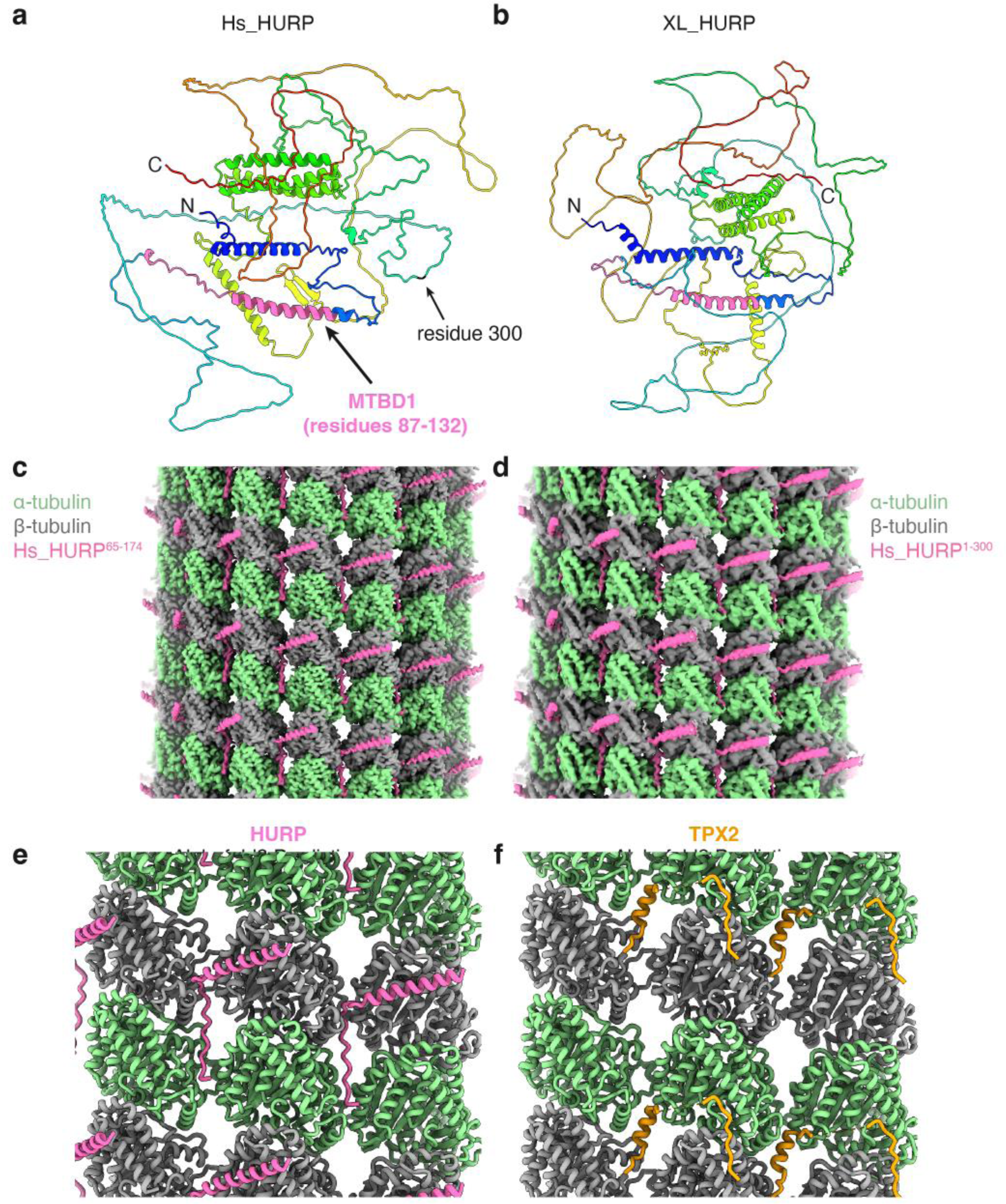
Cryo-EM structure of human HURP constructs on GMPCCP-stabilized lattice. **a**Alphfold2 prediction of human HURP structure colored in rainbow from N- to C- terminus. The microtubule binding domain 1 (MTBD1) identified by our cryo-EM structure is colored pink. **b** Alphfold2 prediction of Xenopus Laevis HURP structure colored in rainbow. Its putative MTBD1 based on sequence alignment is colored in pink. **c** Cryo-EM structure of HURP^65-174^ bound to GMPCPP-MT at 2.8 Å location resolution. **d** Cryo-EM structure of HURP^1-300^ bound to GMPCPP- MT at 3.8 Å location resolution. **e** Atomic model of HURP bound to GMPCPP-MT. **f** Atomic model of TPX2 bound to GMPCPP-MT, from a previous study (PDB ID: 6BJC)^43^.

The resolved cryo-EM density of HURP corresponds to residues 87-132 (**Fig. 5e**), which largely overlaps with the previously identified MTBD1^39^. The fact that both HURP^65-174^ and HURP^1-300^ constructs produce nearly identical MAP densities is consistent with the previous report that MTBD1 is the constitutive MT lattice binding site^39^. We also purified a HURP^1-69^ construct that covers only the MTBD2. However, this construct showed only weak binding to GMPCPP-MTs in a co-pelleting assay. Therefore, we didn’t proceed with the cryo-EM study.

As revealed by our cryo-EM structure of HURP^65-174^ bound to a GMPCPP-MT (**Fig. 5e**), an N-terminal helix of HURP (residues 82-114) binds the external surface of β-tubulin, while a loop following the helix (residues 115-132) binds deeply at the groove between two neighboring protofilaments and simultaneously interacts with four tubulin subunits. This bipartite MT binding mode of HURP is somewhat similar to TPX2 **(Figs. 5e and 5f**) (see Discussion). Such interaction with the MT lattice may explain our observations that HURP reduces the frequency of catastrophes and increases the number of rescue events (**Figs. 4f and 4g**), as HURP binding could strengthen the lateral interaction between protofilaments, including at the MT plus ends.

## Discussion

In this study, we show that HURP is necessary for RanGTP-induced branching MT nucleation in *Xenopus* egg extract. We uncovered the mechanism through which HURP facilitates MT generation, namely by bridging lateral tubulin subunits within the MT lattice. HURP’s binding mode to the MT lattice helps explain how HURP tunes MT dynamic properties such as frequency of catastrophe and rescue events. Other MAPs that bind between protofilaments, such as TPX2^43^, EB family proteins^44,45^, doublecortin^46^, and CAMSAP^47^ also influence MT dynamic properties to some extent. Interestingly, so far only HURP and TPX2 have been found to possess a bipartite MT binding mode, in that they both have one element anchored onto the ridge of one protofilament, and another element binds deeply at the groove between protofilaments (**Figs. 5e and f**). Based on their binding sites on tubulin, both HURP and TPX2 should be able to laterally bridge two tubulin dimers, either in their soluble tubulin form or in their polymerized state (MTs). It is tempting to speculate that HURP may bridge two soluble tubulin dimers or short protofilaments laterally, therefore promoting the formation of early tubulin nucleation intermediates^48^, which may resemble a curved, GTP-rich, sheet-like structures^49–51^. In the cell, HURP may serve to stabilize lateral tubulin interactions that assemble on γ-TuRC to form the critical nucleus or stabilize the nascent microtubule lattice nucleated by γ-TuRC.

In the presence of TPX2, we found that the MT-stabilizing function of HURP favors branching MT nucleation as opposed to the formation of single microtubules. We showed that this occurs by formation of a TPX2 condensate that recruits HURP as a client protein to the MT surface. Therefore, we propose a model in which TPX2, released by RanGTP, preferentially enriches HURP to pre-existing MTs through formation of a co-condensate to promote MT formation via branching MT nucleation (**Fig. 6**). TPX2 also enriches γ-TuRC, augmin, and soluble tubulin to MTs^7,11^, all necessary for branching MT nucleation. Therefore in this model, TPX2 serves to efficiently localize all components together to facilitate branching MT nucleation during early spindle assembly, with HURP serving as a critical stabilizing protein for daughter MTs.

**Figure 6:**
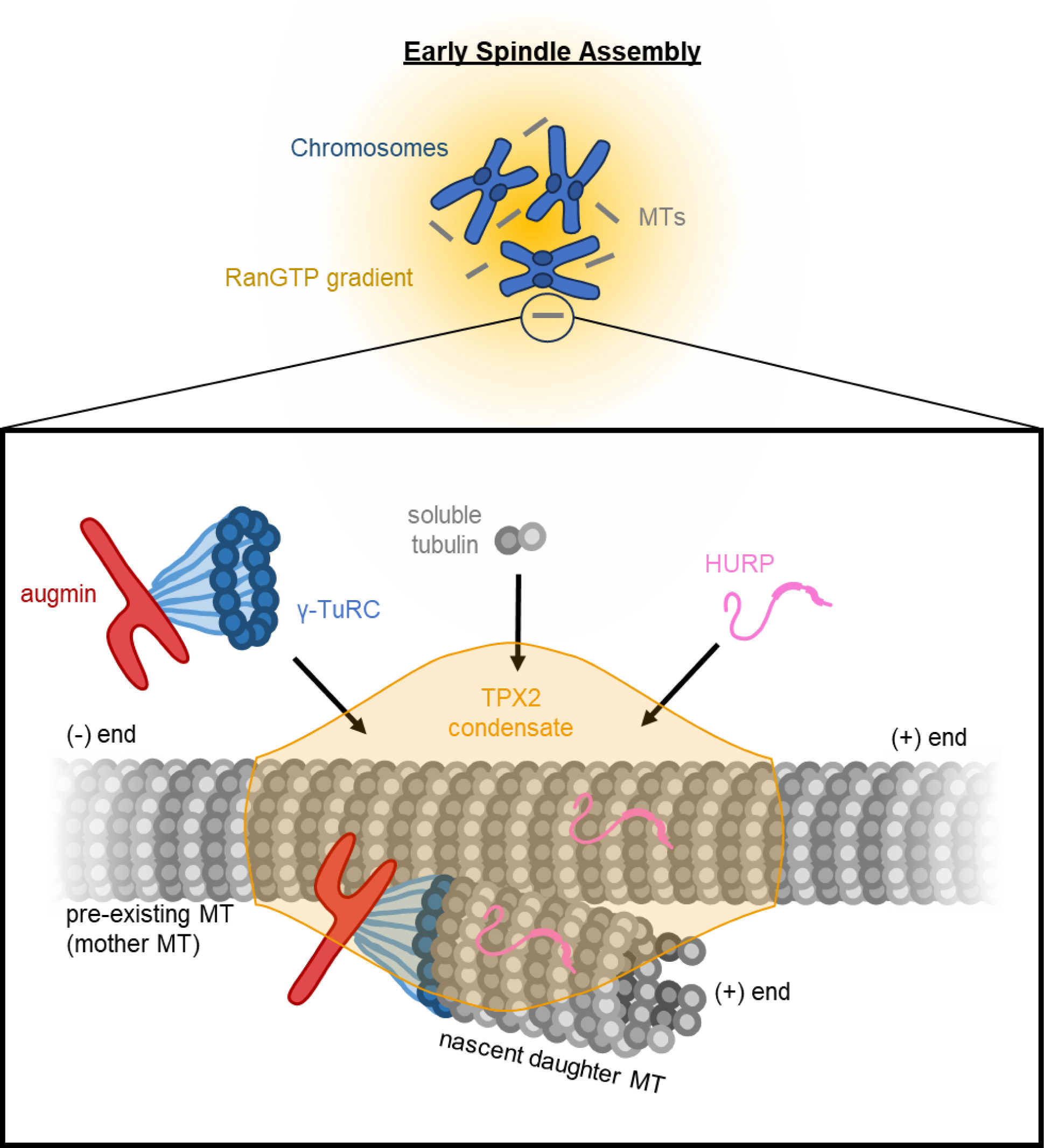
Model for HURP-TPX2 relationship in branching microtubule nucleation. During early spindle assembly, RanGTP produced by chromosomes creates a gradient of RanGTP. RanGTP releases spindle assembly factors augmin, TPX2, and HURP, allowing them to individually bind to forming microtubules. Upon binding to a microtubule, TPX2 forms a condensate and recruits soluble tubulin to the lattice of the pre-existing microtubule. Additionally, the TPX2 condensate enriches for augmin, γ-TuRC, and HURP, forming a site enriched in components necessary for branching microtubule nucleation. Being enriched at this site, HURP will bind and stabilize the pre-existing mother microtubule, as well as the nascent daughter microtubules that form via branching microtubule nucleation.

Apart from RanGTP, other studies have determined that the function of human HURP is also positively regulated by Aurora A phosphorylation. Phosphorylation of the C-terminus of human HURP is necessary for localization to spindle MTs^29,52^. Furthermore, this phosphorylation is hypothesized to contribute to its dynamic localization to the MTs near chromosomes during metaphase^29^. At metaphase, active Aurora A and TPX2 are bound and localized to spindle poles^53^. Given our data, we suggest that if TPX2 is sufficiently concentrated at the poles, TPX2 may condense and enrich HURP, resulting in more efficient phosphorylation of HURP by Aurora A and thus maintenance of HURP localization in the metaphase spindle.

Here, we determined that TPX2 and HURP fulfill independent functions, both crucial to branching MT nucleation, and working synergistically to accomplish one output. We show that unlike TPX2, HURP is incapable of forming condensates or localizing tubulin and γ-TuRC to the MT. And unlike HURP, we have previously shown that TPX2 has little effect on the number of MTs formed from γ-TuRC in vitro^35^. Our results are consistent with previous findings in *Xenopus* egg extract, which determined that TPX2 and HURP cannot compensate for the loss of one another in chromatin-mediated MT formation^27,28^.

However, in *Drosophila* oocytes, TPX2 is dispensable for chromatin-dependent MT generation, while HURP is still required^17^. This deviation is consistent with differences in the *Xenopus* and *Drosophila* proteins. We have previously found that the N-terminal portion of TPX2 drives TPX2 condensation^11^, yet *Drosophila* TPX2 only shares homology with the C-terminal domain^17,54^. Since our model relies on the ability of TPX2 to efficiently form condensates, it is reasonable that the role of TPX2 in *Drosophila* would not be entirely consistent with *Xenopus*. In human cells, the role of TPX2 in branching MT nucleation has yet to be fully studied. However, a recent study revealed that HURP interacts with TPX2 in HeLa cells^29^, suggesting that our model could be conserved in human cells.

Given the localization of HURP to kinetochore fibers^18,28^, our findings explain how HURP may actively contribute to augmin-mediated k-fiber amplification and maintenance during spindle assembly^4^. It was previously suggested that HURP confers k-fiber MT stability through its ability to bundle MTs together^18,28^. In our study, we find that HURP binding to a single MT is sufficient to confer stability that results in an increase of MTs being formed from γ-TuRC in vitro. However, MT bundling (e.g., bundling of a nascent daughter MT to the mother MT) could provide additional stability for MT formation. If true, this would be impactful for k-fiber formation and maintenance, as these MTs are highly bundled.

By adding a microtubule stabilization factor to the proteins that regulate microtubule formation, we bring the microtubule field on par with the actin field. Just like the Arp2/3 complex, formins, and Spire were discovered to seed and bring together actin filaments^55^, their microtubule counterparts during mitosis are now discovered to be γ-TuRC, XMAP215/chTOG, and HURP. The fact that HURP is locally regulated helps explain how it functions specifically in branching microtubule nucleation. We believe this concept helps further our understanding of how the cell can form the microtubule cytoskeleton at the right time and location to enable cell function.

## Material and Methods

### Ethics

*Xenopus laevis* husbandry was done in accordance with the NIH Guide for the Care and Use of Laboratory Animals and the approved Institutional Animal Care and Use Committee (IACUC) protocol 1941 of Princeton University.

### Cloning

The full-length *Xenopus* HURP sequence (NCBI Reference Sequence: XP_018087757.1) was synthesized (Genscript) with codon optimization for expression in *Escherichia coli* in a pUC57 vector. This sequence was inserted into a modified pST50 vector^56^ containing N-terminal Strep-6xHis-TEV-GFP-PreScission tags using Gibson assembly (NEB Cat #: E2611L) to obtain a tagged construct of full-length HURP. The assembled construct was transformed into chemically competent DH5ɑ *Escherichia coli* cells (NEB Cat #: C2987I), spread on selective Luria broth (LB) agar plates containing 50 μg/mL carbenicillin (GoldBio Cat #: C-103-25), and grown in a 37 °C incubator overnight. Single colonies from the selection plates were grown in selective LB media containing 50 μg/mL ampicillin (Goldbio Cat #: A-301-55) overnight in a 37 °C shaking incubator (180 rpm). Plasmids were purified from overnight cultures using the QIAprep Spin MiniPrep Kit (Qiagen Cat #: 27106). Proper assembly of the construct was screened for by restriction digest of the purified plasmids followed by gel electrophoresis. Final confirmation of the construct was done by Sanger sequencing using T7 and T7-Term sequencing primers (Genewiz). An identical strategy was used to create a full-length HURP construct with N-terminal Strep-6xHis-TEV-mCherry-PreScission tags, as well as a Strep-6xHis-TEV tagged construct of the HURP(462-576) fragment used for antibody production.

### Expression and Protein Purification

Both HURP plasmids were transformed into Rosetta2 (DE3) *Escherichia coli* cells (NEB: C29871) and grown in 2 L of LB media containing 50 μg/mL ampicillin and 25 μg/mL chloramphenicol at 37 °C in a shaking incubator (200 rpm). After reaching an OD600 of 0.6, protein expression was induced by addition of 0.5 mM isopropyl-β-D-thiogalactoside (IPTG). The full-length HURP construct was expressed for 4 hours at room temperature in a shaking incubator (200 rpm). The HURP(462-576) fragment was expressed for 16-18 hours at 16 °C in a shaking incubator (200 rpm). After expression, the cultures were pelleted, flash-frozen, and stored at −80 °C.

Purification of the full-length HURP constructs was done by thawing a 2 L cell pellet on ice and resuspended up to a volume of 40 mL in His lysis buffer (50 mM NaH2PO4, pH=8.0, 500 mM NaCl, 20 mM Imidazole, 1 mM MgCl2, 6 mM β-mercaptoethanol (BME), 200 μM phenylmethylsulfonyl fluoride (PMSF), 10 μg/mL DNase I) containing a single dissolved cOmplete EDTA-free Protease Inhibitor Cocktail tablet (Roche: 11873580001). The mixture was first gently homogenized with a Biospec Tissue Tearor (Dremel, Racine, WI). The cell mixture was then lysed by high pressure homogenization in an Emulsiflex C3 (Avestin, Ottawa, Canada) by passing the lysate through the Emulsiflex four times at 10,000-15,000 psi. Cell lysate was spun at 30,000 rpm for 30 minutes at °C in a 45Ti rotor using a Beckman Optima-XE 100 ultracentrifuge. Supernatant was passed through a 5 mL column volume (CV) of Ni-NTA agarose resin (Qiagen: 30250). The column was then washed with 10 CV of His binding buffer (50 mM NaH2PO4, pH=8.0, 500 mM NaCl, 20 mM Imidazole, 1 mM MgCl2, 6 mM BME, 200 μM PMSF). Elution of the protein was eluted by flowing 1.5 CV of His elution buffer (50 mM NaH2PO4, pH=8.0, 500 mM NaCl, 250 mM Imidazole, 1 mM MgCl2, 6 mM BME, 200 μM PMSF) through the resin and collecting the flow-through.

The eluate was further purified using size exclusion chromatography. Eluate was concentrated to a volume of ∼750 μL using a 50 kDa molecular weight cut-off (MWCO) spin concentrator (Millipore Sigma: UFC905024). The concentrated eluate was divided into three, 250 μL samples and individually run through a Superdex 200 Increase 10/300 GL column (Cytiva, Marlborough, MA) hooked up to an ӒKTA Pure system. A 500 μL manual injection loop was used for loading and the run was performed in Cytostatic Factor Extract Buffer (CSF-XB) (10 mM HEPES, pH 7.7 using KOH, 100 mM KCl, 1 mM MgCl2, 0.1 mM CaCl2, 5 mM EGTA, 10% v/v sucrose) with a flow rate of 0.5 mL/min. Eluate was collected in 0.5 mL fractions and absorbance at 280 nm was used to track the protein peak. Yield and purity of fractions were assessed by SDS-PAGE gel and Coomassie stain. Fractions with high yield and purity for the full-length HURP construct (MW: 127 kDa) were pooled, spin-concentrated as described above, and snap-frozen in 5 μL aliquots at ∼15 μM. Protein concentration was determined via densitometry using BSA standards on an SDS- PAGE gel stained with Sypro Protein Gel Stain (ThermoFisher: S12000).

The HURP(462-576) construct used in generation of the HURP antibody was purified by His and Strep affinity chromatography. The cell lysis and His purification is carried out the same as the full-length HURP construct, except that buffers contained 300 mM NaCl instead of 500 mM. Eluate from the His purification was then diluted ∼7-fold with Strep binding buffer (50 mM Tris-HCl, pH 8.0, 300 mM NaCl, 1 mM MgCl2, 6 mM BME, 200 μM PMSF) before flowing over 5 mL CV of Strep-Tactin Superflow resin (Neuromics: 2-1206-025). The column was then washed with 10 CV of Strep binding buffer before being eluted with 1.5 CV of elution buffer (Strep binding buffer + 3.3 mM D-desthiobiotin). Yield and purity of the HURP(462-576) construct (MW: 17 kDa) was assessed by SDS-PAGE gel and Coomassie stain. A concentration of ∼1 mg/mL was determined by Bradford assay.

An existing construct of full-length *Xenopus laevis* TPX2 (Strep-6xHis-GFP-TPX2) was expressed and purified as previously described^11^. Briefly, expression was induced with 0.75 mM IPTG and expressed for 7 hours at 27 °C. The construct was first purified from lysate by His affinity chromatography in a Tris-based buffer, pH 7.75, notably containing 750 nM NaCl. Eluate from the His purification was then further purified by size-exclusion chromatography using a Superdex 200 HiLoad 16/600 column (GE Healthcare: 28-9893-35) in CSF-XB buffer. Peak fractions were then pooled, concentrated by spin centrifugation (Millipore Sigma: UFC905024), flash frozen, and stored at −80 °C.

Octameric augmin complex (HAUS subunits 1-8, with the HAUS6 construct lacking its C-terminus and containing N-terminal ZZtag-PreScission tags, HAUS3 and HAUS8 containing N-terminal Strep-GFP tags, and HAUS2 containing C-terminal GFP-6xHis tags) was expressed and purified from Sf9 cells as described previously^57^. Briefly, the complex was first purified by IgG Sepharose affinity chromatography in a Tris-based buffer (pH 7.7), eluted by PreScission cleavage. Eluate was then spin concentrated and further purified by size-exclusion chromatography using a Superose 6 Increase 10/300 in CSF-XB with 1 mM dithiothreitol (DTT). Purity of peak fractions were analyzed by SDS-PAGE, pooled, flash frozen, and stored in −80 °C.

An existing construct of EB1-mCherry (EB1-6xHis-mCherry in a pET21a backbone) was expressed and purified as previously described previously^58^. Briefly, the protein was purified by His affinity chromatography in sodium phosphate buffers, pH 7.4. Eluate was purified further by size-exclusion chromatography using a Superdex 200 pg 16/600 (GE Healthcare) in CSF-XB. Purity of peak fractions were analyzed by SDS-PAGE, pooled, flash frozen, and stored in −80 °C.

Expression and purification of Ran(Q69L) (Strep-6xHis-TEV-BFP-Ran(Q69L)) was done similarly to a previously published protocol^58^. Specific to this study, expression of the construct was induced with 0.5 mM IPTG after an OD600 of 0.8 was reached in Terrific Broth (TB) media. Expression was carried out for 18 hours at 16 °C. Strep affinity chromatography and subsequent dialysis into CSF-XB was carried out as described previously^58^.

Purification and biotinylation of the native *Xenopus laevis* γ-TuRC complex was done as described previously^36^. Briefly, Halo-Magne beads (Promega: G7287) were coated in purified HaloTag-PreScission-humanγTuNA. The beads were then incubated with thawed *Xenopus laevis* extract to allow for γTuNA to bind to native γ-TuRC. After incubation, beads were washed and incubated with NHS-PEG4-Biotin (ThermoFisher: A39259). After incubation, beads were then resuspended and incubated overnight in a PreScission protease cleavage buffer to allow for cleavage of γTuNA-γTuRC from the HaloTag-bound beads. After cleavage, the supernatant was obtained, concentrated, and spun through a sucrose gradient. Peak fractions from the sucrose gradient were determined by Western blot, then flash frozen and stored at −80 °C. Concentration of γTuRC was assumed to be between 150-200 nM, as previously determined^36^.

Different human HURP constructs covering MTBD1 (residues 65-174), MTBD2 (residues 1-69) and MTBD1-2 (residues 1-300) were cloned into pMBP vector with an N-terminal MBP tag. The recombinant proteins were overexpressed in E. coli BL21(DE3) at 16 °C overnight after induction by 0.2 mM IPTG at OD600 of 0.8. The proteins were purified by Amylose resin (NEB) and eluted with 10 mM maltose. Then the eluate was further purified by gel filtration (Superdex 200, GE Healthcare) in a buffer containing 25 mM Tris (pH 8.0), 150 mM NaCl, 1 mM DTT. The peak fraction was collected and concentrated to ∼5 mg/mL, then stored at −80 °C with aliquots of 20 μL.

### Generation of HURP antibody and purification

A total of ∼10 mg of purified HURP(462-576) at ∼1 mg/mL was sent to the Covance company for inoculation of two rabbits to produce polyclonal antibodies against HURP. Antibody was provided by Covance in the form of antisera. The HURP antibody was then purified by first conjugating 10 mg of HURP(462-576) antigen to 0.5 mL of Affi-gel 15 (Bio-Rad: 1536051) resin (brought up to 10 mL with PBS buffer), rotating overnight at 4 °C. Antigen-coupled beads were then washed sequentially with 0.1 M sodium phosphate buffers (pH 7.2), 0.1 M glycine (pH 2.5), then 0.1 M sodium phosphate buffers of pH 7.2, 11, and 7. After washes, the resin was incubated with 3 mL of antisera, brought up to 10 mL with PBS, rotating overnight at 4 °C. Resin was then washed sequentially with sodium phosphate buffers of pH 7, pH 7 with 0.5 M NaCl, and pH 7 again to remove the salt. Antibody was eluted with 0.1 M glycine, pH 2.5, and quickly neutralized with 1 M HEPES buffer, pH 7.7, at a 1:10 glycine:HEPES ratio. Elution was done in 1 mL fraction volumes (after neutralization with HEPES buffer). Yield was determined by A280 via NanoDrop. Peak fractions were combined, flash frozen in 40 μg aliquots, and stored at −80 °C.

### Tubulin labeling and GMPCPP-stabilized microtubules

Unlabeled cycled tubulin purified from bovine brain was purchased from PurSolutions (Cat#: 032005). Bovine brain tubulin was labeled following previously described methods^59^. Labeling with Cy5-NHS ester (GE Healthcare, PA15101) yielded 54–70% labeling efficiency. Labeling efficiency with Alexa-568 NHS ester (Invitrogen: A20003) was 36–40%. Labeling efficiency with ATTO647N (ATTO-TEC, AD 647N) was 62%. Labeling efficiency with biotin-PEG4-NHS (Thermo Scientific: A39259) was not calculated.

Fluorescent and biotinylated double-cycled GMPCPP-stabilized MTs were made similarly to a previously published protocol^33^. Tubulin was polymerized (40 μL reaction) at a total tubulin concentration of 20 μM in BRB80 buffer (80 mM K-PIPES at pH 6.8, 1 mM EGTA, 1 mM MgCl2) containing 1 mM GMPCPP. Of the 20 μM total tubulin, 2 μM is composed of ATTO647N-labeled tubulin, 2 μM is composed of biotin-labeled tubulin, and the rest is composed of unlabeled tubulin. The reaction was assembled over ice. Polymerization was performed in 37 °C water bath for 30 min, protecting the reaction from light. MTs were then pelleted by centrifugation in a tabletop centrifuge at 13,000 rpm for 15 minutes (room temp). MTs were then depolymerized by resuspending and incubating the pelleted MTs in ice-cold BRB80 buffer for 20 minutes in a volume totaling 80% of the original volume. GMPCPP was then added to 1 mM, and MTs were again polymerized and pelleted as before. This time, MTs were resuspended in warm BRB80 buffer in a volume totaling 80% of the volume prior to pelleting. MTs were flash frozen in 2 μL aliquots and quickly thawed before use. MTs were diluted 1:100 to 1:1000 prior to use in experiments.

### Immunodepletion of *Xenopus laevis* egg extract and western blot analysis

Meiotic *Xenopus laevis* extract was performed as previously described^60^ and assayed for quality (described below in “TIRF imaging of MT formation in *Xenopus laevis* egg extract”). All steps for the immunodepletion were done on ice and in a 4 °C cold room. For immunodepletions, 300 μL of Dynabeads protein A slurry (Thermofisher: 10001D) was washed thrice 600 μL TBS-T buffer. A magnetic strip was used to separate the beads from the solution for washes. Half of the beads were then resuspended in a HURP antibody solution for HURP depletion. The other half were resuspended in an IgG antibody solution for the mock depletion. Antibody solutions consisted of 40 μg of antibody in a total volume of 180 μL using Tris-buffered saline with Tween-20 (TBS-T) buffer (50 mM Tris, pH 8.0, 138 mM NaCl, 2.7 mM KCl, 0.1% w/v Tween-20). Antibody-bead mixtures were incubated overnight at 4 °C on a rotator. After incubation, beads were washed twice with 600 μL TBS-T and thrice with 600 μL CSF-XB. Both sets of beads were split into three tubes for three rounds of immunodepletion, keeping in CSF-XB. To immunodeplete the extract, consecutive rounds of immunodepletion were performed by first aspirating the CSF-XB from the beads and resuspending gently in 60 μL of prepared extract using a wide bore 100 μL pipette tip. The mixture was incubated on ice for 60 minutes, resuspending gently every 20 minutes with a fresh wide bore pipette tip. After 60 minutes, the process was repeated with the remaining tubes. At the end of the third depletion, extract was collected and placed in a fresh tube to be used for experiments.

For Western blotting, extract samples of the undepleted (input), mock-depleted, and HURP-depleted extract were collected (2 μL of extract in 48 μL 1X SDS Sample Loading Buffer + 50 mM DTT), boiled, and 15 μL of each sample was run on two separate 4–12% Bis-TRIS SDS-PAGE gels (ThermoFisher: NP0321BOX). Proteins were then transferred to a nitrocellulose membrane using the Invitrogen iBlot 2 transfer device (ThermoFisher: IB21001), blocked for 1 hour at room temperature, and cut with a razor blade to allow for probing of multiple factors by Western blot. All probing and blocking of membranes was done in TBS-T + 10% w/v non-fat dairy milk. One membrane was used to probe for HURP (HURP antibody described in this paper, 4 μg/mL) and γ-tubulin (Sigma: T6557, 1:1000). The second membrane was used to probe for XMAP215 (Abcam: ab86073, 1:2000), TPX2 (custom antibody described in Alfaro Aco et al 2017, 4 μg/mL), HAUS1 (custom antibody described in Song et al 2018, 3.6 μg/mL), and α-tubulin (Invitrogen: 62204, 1:1000). Membranes were incubated in primary antibody overnight at 4 °C on a shaker, washed thrice with TBS-T on a shaker over the course of 45 minutes, and then incubated for 1 hour at room temperature with either Mouse-IgG, HRP linked secondary antibody (Amersham: NA931-1ML, 1:2000) or Rabbit-IgG, HRP linked secondary antibody (Amersham: NA934-1ML, 1:2000). Membranes were then washed thrice with TBS-T as before, and then incubated with Amersham ECL Western Blotting Detection Reagents (Cytiva: RPN2232) for chemiluminescence detection. Images were taken using the iBright Imaging system, ensuring that no bands were overexposed. The γ-tubulin membrane was stripped with Stripping Buffer (ThermoFisher:46430), reblocked, and probed for the loading control α-tubulin as described above.

To analyze the Western blot for depletion of factors, images were loaded into the Fiji software^61^ (ImageJ; RRID:SCR_002285) and converted to 8-bit (0-255 intensity range) for ease of processing. The mean intensity of each band was measured by selecting a box that encapsulated each band. Boxes were kept an equal size for each measurement. Additionally, the background mean intensity was also measured for each band by selecting a nearby region of the membrane just above or below the band, again keeping the box size consistent. The mean intensity for each band was subtracted from the background mean intensity and standardized to the respective loading control (α-tubulin band) to account for errors in loading the gel. After standardization, the intensities of the mock- and HURP-depleted bands for each blotted protein of interest were used to calculate percent intensity relative to the input lane. For Fig. 1a, the proportion of intensity relative to the input lane (scaled 0-1) was averaged across the four biological replicates and plotted. Values less than 0 and greater than 1 were converted to 0 and 1, respectively, for plotting and statistical analysis. Statistical significance was determined by unpaired two-tailed t-tests comparing the mock- and HURP-depleted conditions for each protein. Plotting and statistics were performed using GraphPad Prism.

### TIRF imaging of microtubule formation in *Xenopus laevis* egg extract

Meiotic *Xenopus laevis* egg extracts were prepared as previously described^60^. Extracts were either used immediately for experiments or subjected to immunodepletion prior to experiments. Quality of every prepared extract was ensured by performing the branching assay (with Ran(Q69L)), described in detail below, and a negative control of the branching assay (no Ran(Q69L)). A quality extract was defined as forming characteristically dense fan-like structures in the presence of Ran(Q69L) and forming few MTs (<10 MTs in 10 minutes) when no Ran(Q69L) is added.

For branching MT nucleation assays, 7.5 µL of extract was incubated on ice with 0.5 µL of 10 mM vanadate (0.5 mM final), 0.5 µL of 1 mg/mL end-binding 1 (EB1)-mCherry protein (0.05 mg/mL final), 0.5 µL of 1 mg/mL Cy5-tubulin (0.05 mg/mL final), 0.5 µL of 200 μM Ran(Q69L) (10 μM final), and 0.5 µL of CSF-XB. For addback of purified HURP to HURP-depleted extract, CSF-XB was replaced with 0.5 µL of 5 μM purified HURP (250 nM final). If necessary, purified proteins were previously diluted in CSF-XB to obtain the stock concentrations listed.

For assaying the effect of excess HURP on the number of MTs, 7.5 µL of extract was incubated on ice with 0.5 µL of 10 mM vanadate (0.5 mM final), 0.5 µL 1 mg/mL end-binding 1 (EB1)- mCherry protein (0.05 mg/mL final), 0.5 µL of 1 mg/mL Cy5-tubulin (0.05 mg/mL final), 0.5 µL of CSF-XB, and 0.5 µL of purified HURP. HURP protein was diluted in CSF-XB prior to addition into the reaction mix, such that the final concentrations in the reaction mix were as stated in Figs. 2e and 2c. A similar approach was used for the addition of 50 nM TPX2 to the extract for Fig. 2c. For the negative control, addition of HURP was replaced with additional CSF-XB. For reactions containing both HURP and TPX2, all CSF-XB was omitted from the reaction mix and was replaced with 0.5 µL of 2.5 μM HURP (125 nM final) and 0.5 µL of 1 μM TPX2 (50 nM final).

All extract reactions (10 µL total) were gently mixed by pipetting twice with a wide-bore 100 µL pipette tip before adding to a flow-channel at 18-20 °C. Flow channels were assembled by evenly spacing thin strips of double sided tape onto a microscope slide and gently pressing a 20 mm x 20 mm coverslip on top of the tape. In this manner, 2-3 flow channels were created to support the number of reactions being imaged. Reactions loaded in multi-flow channel slides (different conditions) were continuously and sequentially imaged for ∼30 minutes. Time = 0 is defined as the moment the reaction was loaded into the flow-channel.

Multi-channel fluorescence images for all experiments were acquired with no delay between acquisitions using the NIS-Elements AR program (NIKON, ver. 5.02.01-Build 1270; RRID:SCR_014329). The 647 nm/Cy5 channel (excitation: 678 nm, emission: 694 nm) was used for imaging MTs and the 561 nm channel for EB1 localized to MT plus-tips (ex: 587 nm, em: 610 nm). The images were captured on a Nikon Ti-E inverted system (RRID:SCR_021242), with an Apo TIRF 100 x oil objective (NA = 1.49), and an Andor Neo Zyla (VSC-04209) camera with 2×2 binning. Exposure settings for all channels were kept consistent across conditions. Resulting images were 164.92 µm x 139.15 µm. After time-lapse acquisition, a large 3 x 3 multi-channel fluorescence image was acquired to ensure that all time-lapse FOV were representative of the reaction. Stitched 3 x 3 images are provided as supplementary figures for each extract reaction provided. 3 x 3 images were 491.55 µm x 414.75 µm. For all the extract images displayed, the 647 nm channel was pseudo-colored red and the 561 nm channel was pseudo-colored green. Dimensions of representative cropped images are listed in the respective figure legends.

### Analysis of microtubule formation over time in *Xenopus laevis* egg extract

To quantify MT nucleation levels, we extracted the number of MT plus ends (tracked by EB1 spots) for each condition using the 561 nm/EB1-mCherry channel. For early timepoints, the number of MTs were recorded manually in the Fiji software at increments of every 10 frames (∼80 seconds). Manual counting of early timepoints was done due to the inaccuracy of the automated counting system given the inaccurate auto-thresholding when few EB1 foci were present. For later timepoints, we wrote a macro in Fiji to automate counting EB1 spots. Briefly, 110 μm x 93 µm representative windows were cropped from each field of view for the EB1 channel image stack. The image stack was duplicated twice and the Gaussian Blur function was applied to each (one with a sigma(radius) of 1, the other with 5). The 5 radius blur image was subtracted from the 1 radius blur image using the Image Calculator. The subtracted image stack was then autothresholded using the default option, and ‘Analyze Particles’ was used to count the number of thresholded EB1 foci (minimum size = 0.1 μm^2^). Manual and auto-counts were combined and plotted over time (based on the framerate and time imaging began) using GraphPad Prism version 10.0.3 for Windows (GraphPad Software, www.graphpad.com). Each experiment was performed at least three times. Each replicate was performed on separate days, using fresh egg extract prepared from different frogs. All replicates had similar results. No statistics were performed. Representative images are shown in Figs. 1 and 2.

### Preparation of functionalized coverslips and imaging chambers

Functionalized glass coverslips used in all in vitro γ-TuRC nucleation and MT nucleation assays were generated using a previously published method^36^. Briefly, coverslips were cleaned by sonication in a series of 3 M NaOH followed by Piranha solution (2:3 rate of 30% w/w H2O2 to sulfuric acid), rinsed with milliQ water, dried, then treated with 3-glycidyloxypropyl trimethoxysilane (Sigma: 440167) at 75 °C for 30 min. Following treatment, coverslips were washed with acetone, dried with nitrogen gas, and functionalized with a 9:1 mixture of HO-PEG- NH2 to biotin-CONH-PEG-NH2 by weight (Rapp Polymere: 103000–20 and 133000-25-20) at 75°C overnight. After incubation, coverslips were washed with MilliQ water, dried, and stored at 4°C for up to 3 months.

For the γ-TuRC nucleation assays, imaging chambers were made by making a channel on a glass slide with double-sided tape, coating the channel with 2 mg/mL of PLL-g-PEG (in MilliQ water), letting air dry, rinsing with MilliQ water, then drying with nitrogen gas. Functionalized coverslips were then placed on the double-sided tape (functionalized side facing down). Chambers were used within 24 hours. Imaging chambers for the MT localization assays were made similarly except that multi-flow channel slides (2-3 flow channels) were made with the double-sided tape, not coated with PLL-g-PEG, and used within one week stored at 4 °C.

### In vitro γ-TURC-mediated microtubule nucleation assay

Imaging chambers were blocked with room temperature 5% w/v Pluronic F-127, followed by cold assay buffer (BRB80 with 30 mM KCl, 0.075% w/v methylcellulose 4000 cp, 1% w/v D-glucose, 0.02% w/v Brij-35, and 5 mM BME). If performing the γ-TuRC nucleation assay, the assay mix was also supplemented with 1 mM GTP. After blocking for 5 minutes, the imaging chambers were washed with cold 0.05 mg/ml κ-casein in assay buffer, then incubated with cold 0.05 mg/mL NeutrAvidin (ThermoFisher: A2666) on a cold block for 1.5 minutes, then washed with cold BRB80 buffer.

A nucleation mix containing 12.5 μM bovine tubulin (5% AlexaFluor 568-tubulin and the remaining unlabeled) and 1 mg/mL BSA (Sigma: A7906) in cold assay buffer was centrifuged at 80,000 rpm in TLA100 rotor (Beckman Coulter) for 12 minutes at 2 °C (20% loss of tubulin assumed, final tubulin concentration = 10 μM). During the spin, an aliquot of biotinylated γ-TuRC was diluted 1/100 in cold BRB80 buffer, flowed into the imaging chamber, and incubated for 7 minutes at room temperature in a dark humidity chamber. One diluted aliquot was used for all experiments in a given day. After centrifugation of the nucleation mix, glucose oxidase (SERVA GmbH: SE22778, final concentration of 0.68 mg/mL), catalase (Sigma: SRE0041, final concentration of 0.16 mg/mL), and either GFP-HURP (final concentration of 50 nM, pre-diluted in CSF-XB; n=4) or an equal volume of CSF-XB (n=3) was added to the nucleation mix. The imaging chamber was then washed with cold BRB80 and the nucleation mix was loaded into the imaging chamber (t=0) and imaged for 5 minutes (1 frame per 2-3 seconds), starting at t=60 sec. We used the same imaging set-up as our previous work^35,36^, notably a Nikon Ti-E inverted stand (RRID:SCR_021242) with an Apo TIRF 100 x oil objective (NA = 1.49) and an Andor iXon DU- 897 EM-CCD camera with EM gain set to 300. The 561 nm channel was used to image Alexa Fluor 568-tubulin and the 488 nm channel was used to visualize GFP-HURP. Exposure settings for all channels were kept consistent across conditions. We used an objective heater collar (Bioptechs: model 150819–13) to maintain 33.5 °C for our experiments. Resulting images were 81.45 μm x 81.45 μm. Dimensions of representative cropped images in figures are listed in the figure legends.

### Analysis of in vitro γ-TuRC-mediated microtubule nucleation assay data

For each movie, we started by processing the entire field of view (81.45 μm x 81.45 μm) for the 5 minute movies using Fiji. The movies were corrected for thermal drift (translation) by using the StackReg plugin^62^ (RRID:SCR_003070). To correct for severe drift, we cropped the corrected stack to a 75.09 μm x 75.09 μm square, allowing us to perform the StackReg function again. We wrote a macro in Fiji to semi-automate data analysis. The macro generates kymographs (time-space) plots based on manual tracing of individual MTs, and prompts the user to extract relevant parameters from the kymographs.

For each kymograph produced, the user is first prompted to determine if the MT is spontaneously nucleated or γ-TuRC mediated. MT kymographs where the tubulin signal formed a right triangle were considered γ-TuRC-mediated. Random appearances of a long MT or a MT that grew from both ends, indicating that the minus end is not capped by γ-TuRC, were marked spontaneous. The location of each MT was recorded and numbered on a separate Z-projected map to prevent double-counting. If a MT was determined to be γ-TuRC-mediated, we proceeded collecting measurements. We manually recorded the nucleation point (origin) for each MT to determine. We drew a line along the growing edge, extracting the slope to generate the growth speed for that MT (based on frame rate for each movie). If the plus-end was visible for the entire kymograph (minimal MT pivoting from the point of origin, i.e., bound γ-TuRC), we manually determined the number of MT catastrophe and rescue events. Catastrophe was defined as a depolymerization event of at least 3 pixels (18.86 μm) and rescue was defined as MT polymerization of at least 3 pixels (18.86 μm) following a catastrophe event. We drew a line across the growing edge, subtracting time depolymerizing in order to measure total MT polymerization time. In summary, for each MT analyzed, the time of MT nucleation (sec), polymerization rate (μm/min), number of catastrophes, number of rescues, and total polymerization time (sec) was gathered.

Time of nucleation for each MT was analyzed using a Python script in Jupyter Notebook^63^ and averaged across all reactions for each condition. The mean and standard error of the mean (SEM) for the number of MTs over time were plotted as shown in Fig. 4b. To determine the rate of γ- TuRC-mediated MTs formed, we fit the MT number curve for each condition to the following condition as done previously^36^:

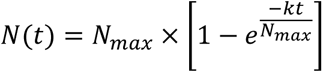

In the equation, *N*(*t*)= the number of MTs nucleated at a given timepoint (t), *N*_*max*_= maximum number of MTs at the final timepoint (360 sec), and *k*= the rate of MT formation. Fitting was done using the scipy.optimize curve_fit function^64^ to determine the *N*_*max*_ and *k*. The *R*^2^ of the fit for each condition are provided in Fig. 4b. Growth rates of all MTs across replicates for each condition were represented in a Tukey box plot using GraphPad Prism. For each replicate in a condition, the average growth rate was obtained and averaged over replicates to obtain a new average and SEM. These combined averages and SEM were then used for statistical analysis by Welch’s two-tailed t-test in GraphPad Prism to determine if the conditions were statistically different. For analysis of catastrophe frequency, we determined how many MTs underwent a catastrophe event versus not undergoing a catastrophe event. Multiple catastrophe events in a single MT were only counted as one. This was done as opposed to obtaining a catastrophe rate (# of catastrophes/total polymerization time) because in many cases for both conditions, MTs did not undergo any catastrophe events in the 360 seconds they were imaged. In addition, we counted how many catastrophe events were rescued. Numbers were pooled across replicates for statistical analysis. The proportions for catastrophe and rescue were plotted and statistically analyzed by Fisher’s exact test in GraphPad Prism to determine if the conditions were statistically different.

### Microtubule localization assays

Imaging chambers were prepped the same as in the first paragraph of the “in vitro γ-TuRC nucleation assay” section. After the NeutrAvidin incubation and cold BRB80 wash, the localization assays were performed as follows:

For the MT localization assay, fluorescent and biotinylated GMPCPP-stabilized MTs were quickly thawed and diluted 1:100 to 1:1000 in warm BRB80, flowed into the imaging chamber(s), and incubated for 7 minutes at room temperature in a dark humidity chamber. A fresh dilution of MTs was used for each slide. ATTO647N-labeled MTs were used for all MT recruitment assays, except for the γ-TuRC recruitment assay which utilized Alexa Fluor 568-labeled MTs. After incubation/binding of MTs in the imaging chamber, all following incubations were performed at room temperature in a dark humidity chamber. MT localization assays were performed as follows:

For recruitment of γ-TuRC to the MT, an estimated ∼70 nM of purified γ-TuRC (assuming 175 nM concentration for undiluted γ-TuRC) was pre-incubated on ice with either 150 nM GFP-augmin, 100 nM GFP-HURP, or an equal volume of CSF-XB for 5 minutes. After incubation of imaging chambers with MTs as stated above, the imaging chambers were washed with 0.05 mg/ml κ- casein in cold assay buffer, then incubated with the pre-incubated protein mix for 5 minutes. After incubation, the chambers were washed with BRB80 buffer and then incubated with 2 μg/mL Alexa Fluor 647-conjugated γ-tubulin (XenC) antibody (described previously^58^) for 10 minutes. After incubation, unbound protein was washed away with cold BRB80 buffer supplemented with 0.68 mg/ml glucose oxidase and 0.16 mg/ml glucose oxidase. Reactions were imaged immediately. For recruitment of soluble tubulin to the MT, 100 nM of Alexa Fluor 568-tubulin was pre-incubated on ice with either 100 nM GFP-TPX2 or 100 nM GFP-HURP for 5 minutes. After incubation of imaging chambers with MTs as stated above, the imaging chambers were washed with 0.05 mg/mL κ-casein in cold assay buffer, incubated with the pre-incubated protein mix for 5 minutes. After incubation, unbound protein was washed away with cold BRB80 buffer supplemented with 0.68 mg/mL glucose oxidase and 0.16 mg/mL glucose oxidase. Reactions were imaged immediately.

For sequential binding reactions, MT-containing imaging chambers were washed with 0.05 mg/ml κ-casein in cold assay buffer, then the first protein (or CSF-XB for the negative control) was flowed in (diluted in CSF-XB to the concentrations indicated in each figure) and incubated for 5 minutes. Unbound proteins were washed with BRB80 buffer then the second protein was flowed in and incubated for 5 additional minutes. After incubation, unbound protein was washed away with cold BRB80 buffer supplemented with 0.68 mg/mL glucose oxidase and 0.16 mg/mL glucose oxidase. Reactions were imaged immediately.

Multi-channel fluorescence images for all experiments were acquired using the NIS-Elements AR program (NIKON, ver. 5.02.01-Build 1270; RRID:SCR_014329). For each assay, a 3 x 3 field of view image was acquired to obtain more MTs for analysis. The 647 nm channel was used for imaging the Alexa647-XenC antibody or ATTO647N-labeled MTs. The 561 nm channel was used to visualize mCherry-HURP, Alexa Fluor 568-tubulin, or Alexa Fluor 568-labeled MTs. The 488 nm channel was used to visualize GFP-HURP, GFP-TPX2, or GFP-augmin. Within each experiment, laser power and exposure settings were kept consistent across conditions, except in Supplementary Fig. S5 where exposure differences are specified in the figure legend. The images were captured on a Nikon Ti-E inverted system (RRID:SCR_021242), with an Apo TIRF 100 x oil objective (NA = 1.49), and an Andor Neo Zyla (VSC-04209) camera with 2×2 binning. Resulting 3 x 3 stitched images were 491.55 µm x 414.75 µm.

### Analysis of microtubule binding/recruitment assays

3 x 3 images were analyzed in Fiji by first isolating the MT channel, auto-thresholding the image using the default setting, and then performing the ‘Open’ and ‘Dilate’ functions sequentially to remove background. ‘Analyze Particles’ of particles greater than 100 pixel units (1.66 μm^2^) was applied and particles added to the ROI manager. The ROIs were then used to measure the intensity of each individual ROI for the MT and protein of interest. Any ROIs associated with an aggregate were excluded. The inverse ROI was used to measure the average background signal. For the inverse ROI, any protein aggregates or isolated phase condensates were excluded from the measurement. Intensity measurements were imported into a spreadsheet, corrected by subtracting the background intensity, and the protein:MT intensity ratios calculated. The average protein:MT intensity for each replicate was calculated and averaged within each condition. Averages for each condition in a given experiment were plotted on a bar chart and statistically analyzed by Welch’s two-tailed t-test using GraphPad Prism to determine if conditions were statistically different. All MT binding assays for any given experiment were either done on the same day or on two separate days. Reaction mixes for replicates performed on the same day were prepared independently. Purified proteins used for all replicates in any given experiment were derived from a single protein prep. Exact number of replicates for all figures are notated in the respective figure legend.

### Bulk phase and co-condensate assay

Purified GFP-TPX2 and ATTO647N-labeled tubulin stocks were precleared of aggregates by centrifugation at 80,000 RPM for 10 min at 4 °C in a TLA-100 rotor. The supernatants were collected and kept on ice. A flow chamber was then washed with 40 μL of cold assay buffer (BRB80 + 50 μg/mL κ-casein). Purified GFP-TPX2, ATTO647N-labeled tubulin, and mCherry-HURP were diluted to their target concentrations in assay buffer, mixed thoroughly, and pipetted into the flow chamber. The flow chamber was sealed with nail polish and incubated coverslip side down for 10 min to allow condensates to settle. Condensates were imaged in bulk and on the coverslip surface using epifluorescence on a Nikon Ti-E microscope with a 100x objective and 1.49 numerical aperture. Exposure times and LED power were consistent across all tested concentrations for each protein. An ORCA-Fusion BT digital CMOS camera was used for acquisition. Condensation was determined visually by assessing whether each condition promoted the formation of mesoscale droplets.

### Cryo-EM sample preparation

GMPCPP-MTs were prepared using an established protocol^44^. Polymerized GMPCPP-MTs were diluted to 0.1 mg/mL concentration and applied to a glow-discharged C-flat 1.2/1.3-4C holey carbon EM grid (Protochips) and incubated inside a Vitrobot (Thermo Fisher Scientific) set at 25°C and 95% humidity for 1 min. Then the grid was washed twice with 3 μL of room temperature HURP proteins (at 1 mg/mL concentration) to allow maximum decoration, before blotting and vitrification in liquid ethane.

### Cryo-EM data collection

Cryo-EM data were collected using a 300 keV Titan Krios microscope equipped with a Cs-corrector (Thermo Fisher Scientific) and a Bioquantum Energy Filter (Gatan) at the Center for Cellular Imaging (WUCCI) in Washington University in St. Louis. In total around 2,000 movies were collected using a K2 Summit direct electron detector (Gatan) in counting mode and an exposure rate of 8.5 electrons/pixel/s on the detector camera. The images were recorded at a nominal magnification of 81,000′, corresponding to a calibrated pixel size of 1.39 Å, with a defocus range from −1.0 to −2.5 μm. A total exposure time of 9 s, corresponding to a total dose of 39.6 electrons/Å2 on the specimen, was fractionated into 30 movie frames. The data were collected automatically using EPU software (Thermo Fisher Scientific). Data collection statistics are reported in **Supplementary Table 1**.

### Cryo-EM data processing

MTs were automatically picked from 1,728 motion-corrected micrographs using template based ‘Filament Tracer’ in CryoSPARC^40^.The templates were a few 2D class averages from a small subset of MTs that were picked manually. The step size between adjacent MT particle boxes was set to be 83 Å, which is the length of an α,β-tubulin heterodimer. Two-dimensional classification was used to remove junk particles and off-centered MT particles (**Supplementary Fig. S8**). In the next step, we performed supervised 3D classification of the MT particles using two reference models (13-PF and 14-PF MTs), and around 80% of the MT particles are 14-PF MTs, which is typical for GMPCPP-MTs. We used only the 14-PF MTs for subsequence data processing in 5 major steps. In step 1, we used ‘Helical Refinement’ function in CryoSPARC to obtain a 3D reconstruction by specifying the 14-start pseudo-helical symmetry (rise 83.0 Å, twist 0°), and then fed the outputs to ‘Local Refinement’ function to further refine the structure with a cylindrical mask covering most of the MT density. In step 2, we used pyem (10.5281/zenodo.3576630) to convert the alignment parameters from CryoSPARC to Relion format, and then to FREALIGN format using customized python scripts. In step 3, we used a previously established protocol to determine the seam location for each MT particle^41^, and imported the alignment parameters with the correct seam location back to CryoSPARC. In step 4, we performed local refinement (using a cylindrical mask covering most of the MT density), local CTF refinement and another round of local refinement, resulting in a MT C1 reconstruction at 3.3 Å resolution. In step 5, to fully utilize the pseudo-helical symmetry of MT, we did symmetry expansion using the 3-start symmetry (rise 8.9 Å, twist −25.8°) and helical order of 11, resulting in 11-fold increase of the total particles number (in theory one can set the helical order to be 14), and performed local refinement of all the symmetry expanded MT particles using a cylindrical mask covering most of the MT density. In step 5, we continued the local refinement using a shorter cylindrical mask (200 Å in length) that covers only 2 protofilaments, resulting in the best local resolution of 2.8 Å.

### Protein identification, model building and refinement

We used the cryo-EM density map of HURP65-174 decorated MT at 2.8 Å local resolution as the input for a deep-learning based software ModelAngelo^42^, which automatically determines the protein sequence for a density and builds tentative atomic models. A search of ModelAngelo derived protein sequence against the entire human proteome produced a single hit of HURP protein with high confidence. The hit was further verified by careful visual inspection of the fit between the atomic model and cryo-EM density map using Coot^65^. The atomic models of α/β- tubulin were adapted from PDB 6DPU^66^, and adjusted in Coot based on the current density map. Atomic model for a patch of 3×3 tubulin dimers were built and refined into the sharpened cryo-EM density map using Phenix.real_space_refine v1.20.1-4487. Secondary structures, Ramachandran, and rotamer restraints were applied during refinement. The quality of the refined model was assessed by MolProbity^67^, with statistics reported in **Supplementary Table 1**.

### Reported statistics

For all results, statistical significance is defined as a p-value ≤ 0.05 and is denoted by a single asterisk (*). P-values ≤ 0.01 were denoted with a double asterisk (**). P-values ≤ 0.001 were denoted with a triple asterisk (***). Not significant results (p-value > 0.05) were denoted with “n.d.”. Statistical methods for determining significance were reported in each individual figure legend and in the methods for each assay.

## Acknowledgements

We especially thank Collin McManus, Jodi Kraus, and Sophie Travis of the Petry Lab for providing reagents for this work and feedback on the manuscript. We thank all former and current members of the Petry and Zhang Lab for any training, discussion, and advice pertaining to this work. V.A.V was supported by NIH Training Grant (T32GM007388) and the Princeton University Charlotte Elizabeth Procter Fellowship. R.Z. is supported by NIGMS grant 1R01GM138854. S.P. is supported by NIGMS grant 1R01GM141100-01A1.

## Author Contributions

V.A.V. conceptualized, designed, performed, and analyzed all MT localization assays, nucleation assays, and experiments in *Xenopus* egg extract. B.G. designed and performed all phase assays. M.M. purified different human HURP constructs and prepared the cryo-EM samples. M.M. and R.Z. collected and processed the cryo-EM data. R.Z. developed the new CryoSPARC-based pipeline for MT data processing. V.A.V. and R.Z. wrote the first draft of the manuscript. All authors contributed to manuscript editing and data interpretation. S.P. and R.Z. supervised the project and acquired funding.

## Data Availability

The single-particle cryoEM structures of HURP^65-174^-decorate GMPCCP-MT has been deposited in the Electron Microscopy Data Bank (EMDB) with accession code EMD-XXXXX. He refined atomic model for a patch (3×3) of tubulin dimers with HURP decoration has been deposited in the Protein Data Bank (PDB) with ID code XXXX.

## Code availability

Python scripts needed for the new CryoSPARC- and FREALIGN-based MT data processing pipeline is available at https://github.com/rui--zhang/Microtubule.

---

**Supplementary Figure 1:**
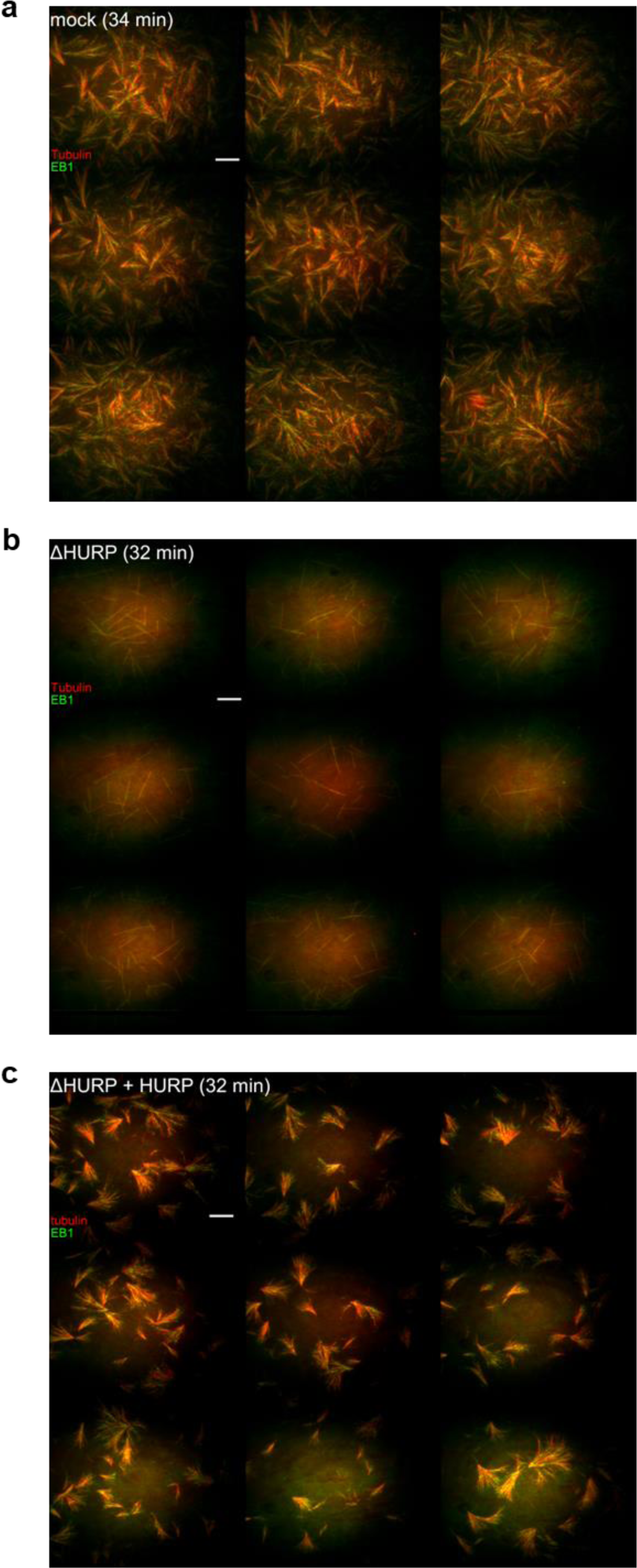
Representative images of branching microtubule nucleation assays. 3 x 3 images taken of the branching microtubule (MT) nucleation reactions after obtaining the movies displayed in Supplementary Movie 1. This was done to ensure that each movie obtained was representative of the reaction **a** 3 x 3 image of the mock depletion condition after 34 minutes **b** 3 x 3 image of HURP-depleted condition after 32 minutes **c** 3 x 3 image of the HURP-depleted + 250 nM purified HURP condition after 32 minutes. MTs (Alexa647-tubulin) are pseudo-colored red and EB1-mCherry is pseudo-colored green for display purposes. Total field of view is 491.55 µm x 414.75 µm. Scale bars = 20µm.

**Supplementary Figure 2:**
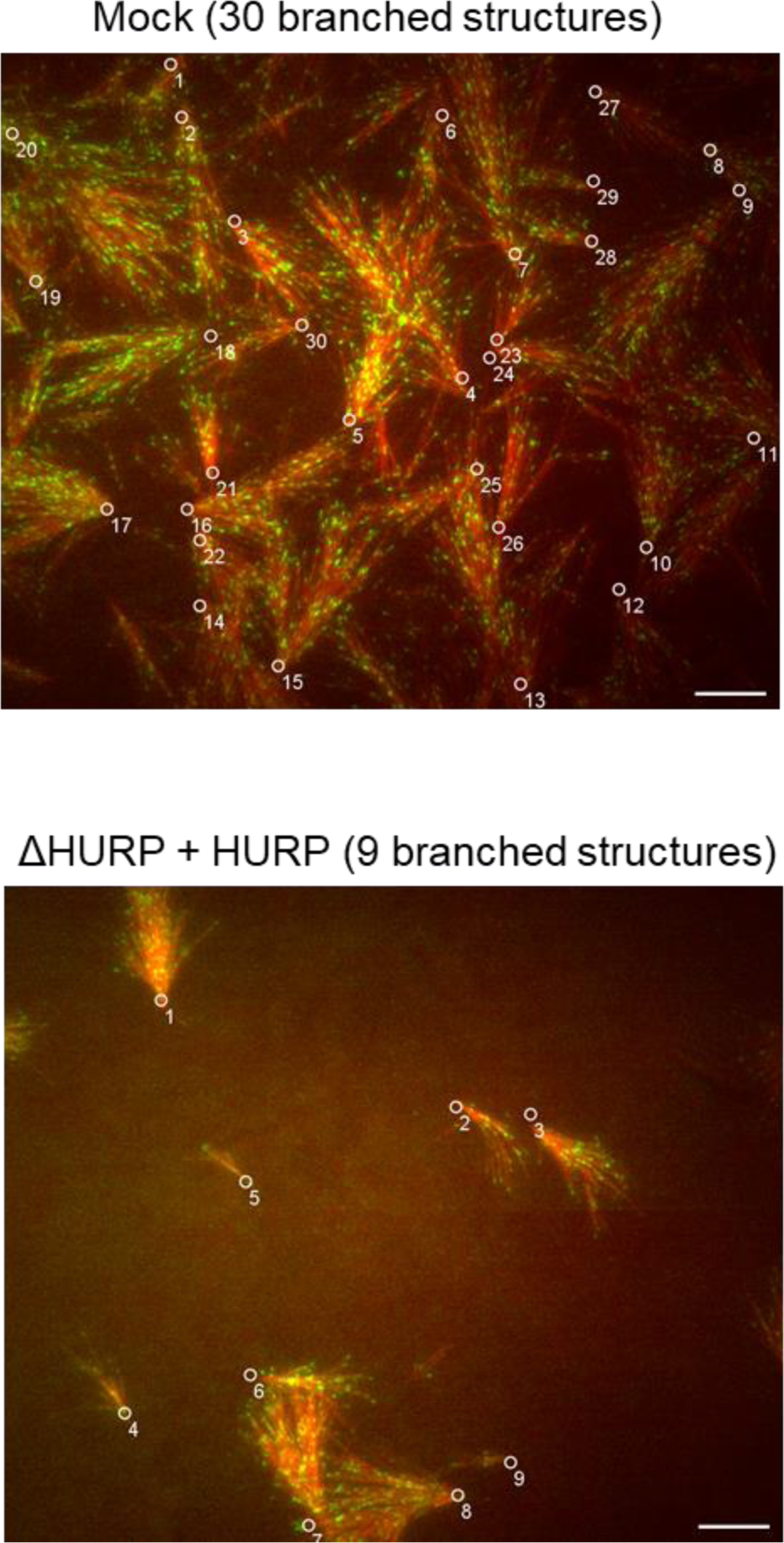
Number of branched structures in mock-depleted and ΔHURP + HURP *Xenopus* egg extract. Representative 110.03 µm x 92.77 µm cropped images used for obtaining the number of microtubules (MTs) over time (EB1 foci per frame) in Supplementary Movie 1 are displayed for the mock and ΔHURP + 250 nM HURP conditions at a timepoint of ∼25 minutes. This timepoint was chosen because individual structures could still be discerned by eye. Individual structures in each condition are denoted with circles and numbered (mock = 30 structures; ΔHURP + HURP = 9 structures). The number of fans was used to normalize the number of MTs over time (EB1 foci per frame) for Fig. 1c. ΔHURP depleted extract was not included because no branched structures were formed. MTs (Alexa647-tubulin) are pseudo-colored red and EB1-mCherry is pseudo-colored green for display purposes. Scale bars = 10 µm.

**Supplementary Figure 3:**
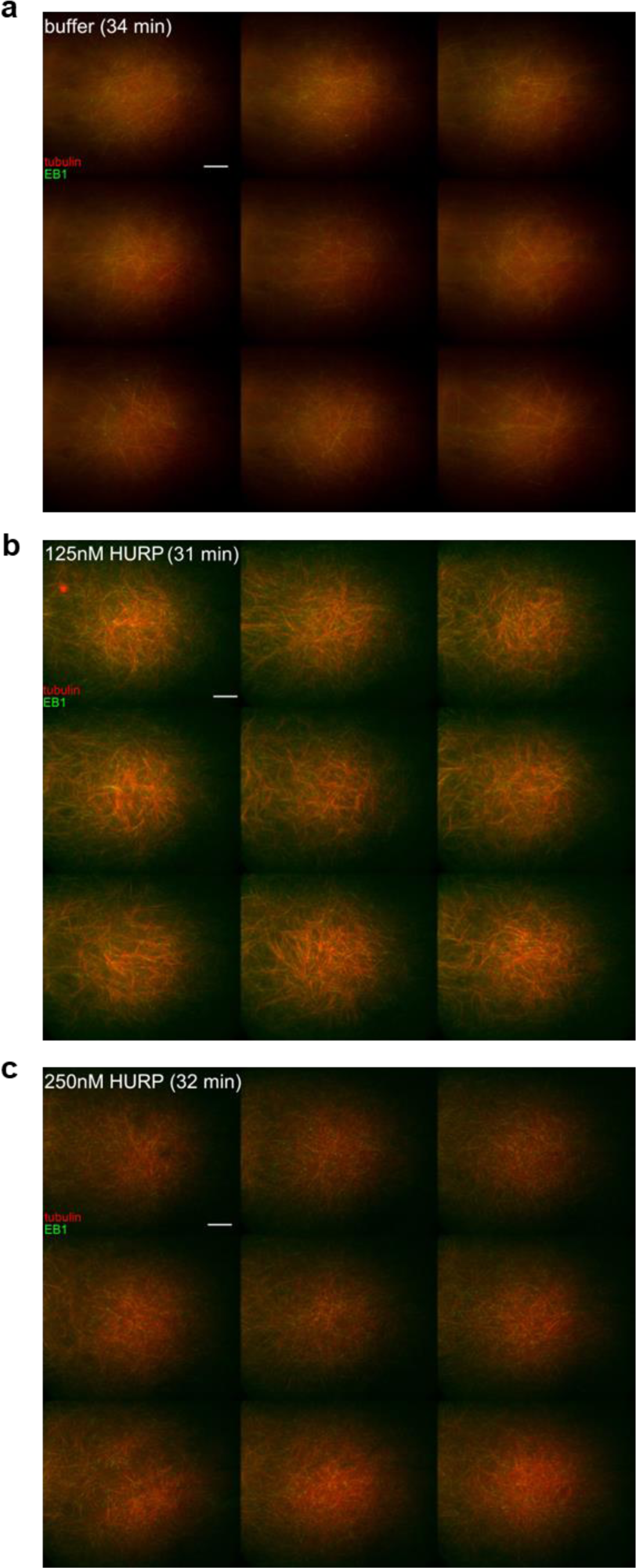
Representative images of HURP-induced microtubule formation in *Xenopus* egg extract. 3 x 3 images taken of the microtubule (MT) nucleation assays with varying concentrations of HURP after obtaining the movies displayed in Supplementary Movie 2. This was done to ensure that each movie obtained was representative of the reaction **a** 3 x 3 image of the buffer condition after 34 minutes **b** 3 x 3 image of the 125 nM HURP after 31 minutes **c** 3 x 3 image of the 250 nM HURP condition after 32 minutes. MTs (Alexa647-tubulin) are pseudo-colored red and EB1- mCherry is pseudo-colored green for display purposes. Total field of view is 491.55 µm x 414.75 µm. Scale bars = 20 µm.

**Supplementary Figure 4:**
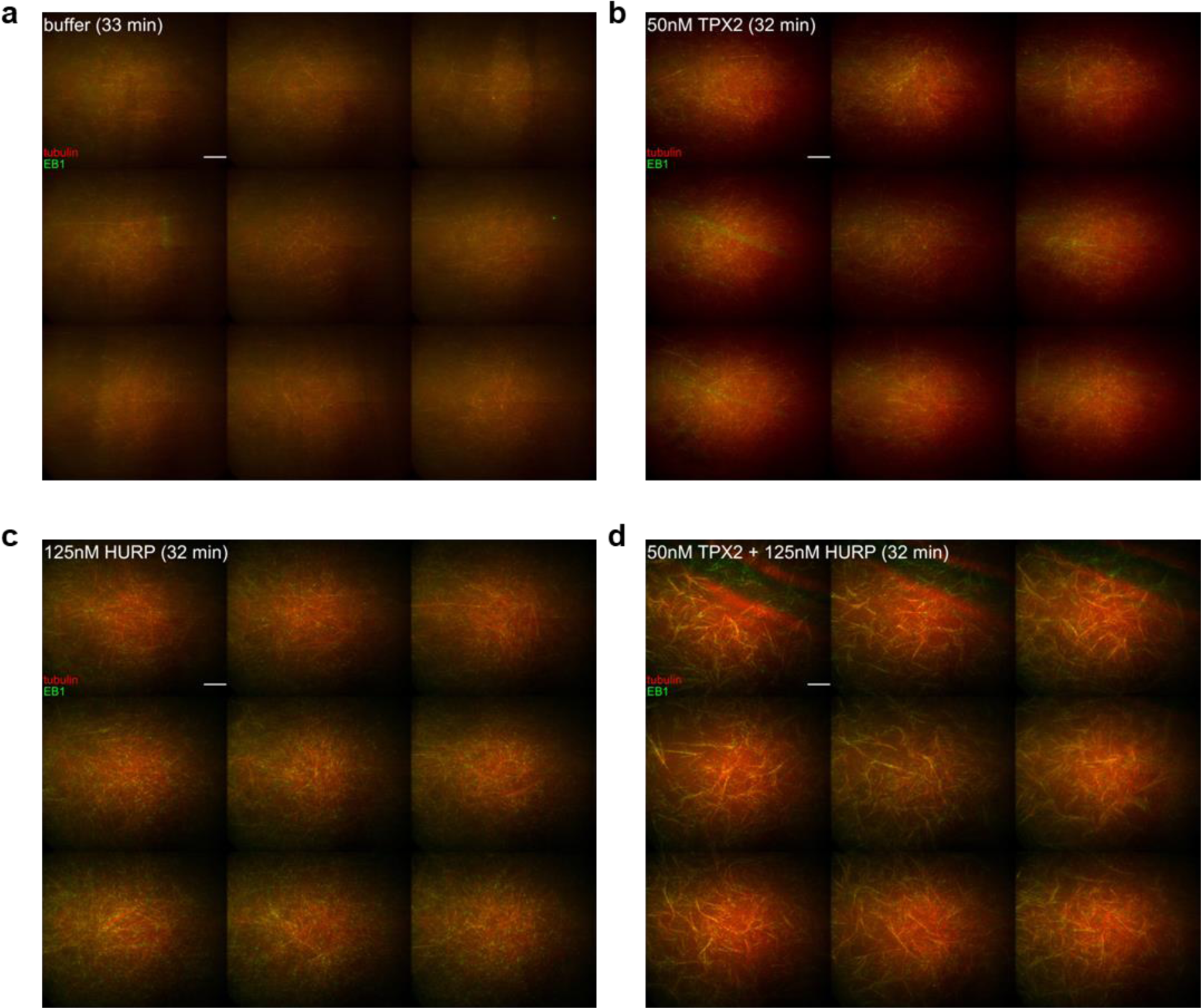
Representative images of TPX2 and HURP synergy in *Xenopus* egg extract. 3 x 3 images taken of the microtubule (MT) nucleation assays with HURP and TPX2 after obtaining the movies displayed in Supplementary 2. This was done to ensure that each movie obtained was representative of the reaction **a** 3 x 3 image of the buffer condition after 33 minutes **b** 3 x 3 image of the 50 nM HURP after 32 minutes **c** 3 x 3 image of the 125 nM HURP condition after 32 minutes. **d** 3 x 3 image of the 50 nM TPX2 + 125 nM HURP condition after 32 minutes. MTs (Alexa647-tubulin) are pseudo-colored red and EB1-mCherry is pseudo-colored green for display purposes. Total field of view is 491.55 µm x 414.75 µm. Scale bars = 20 µm.

**Supplementary Figure 5:**
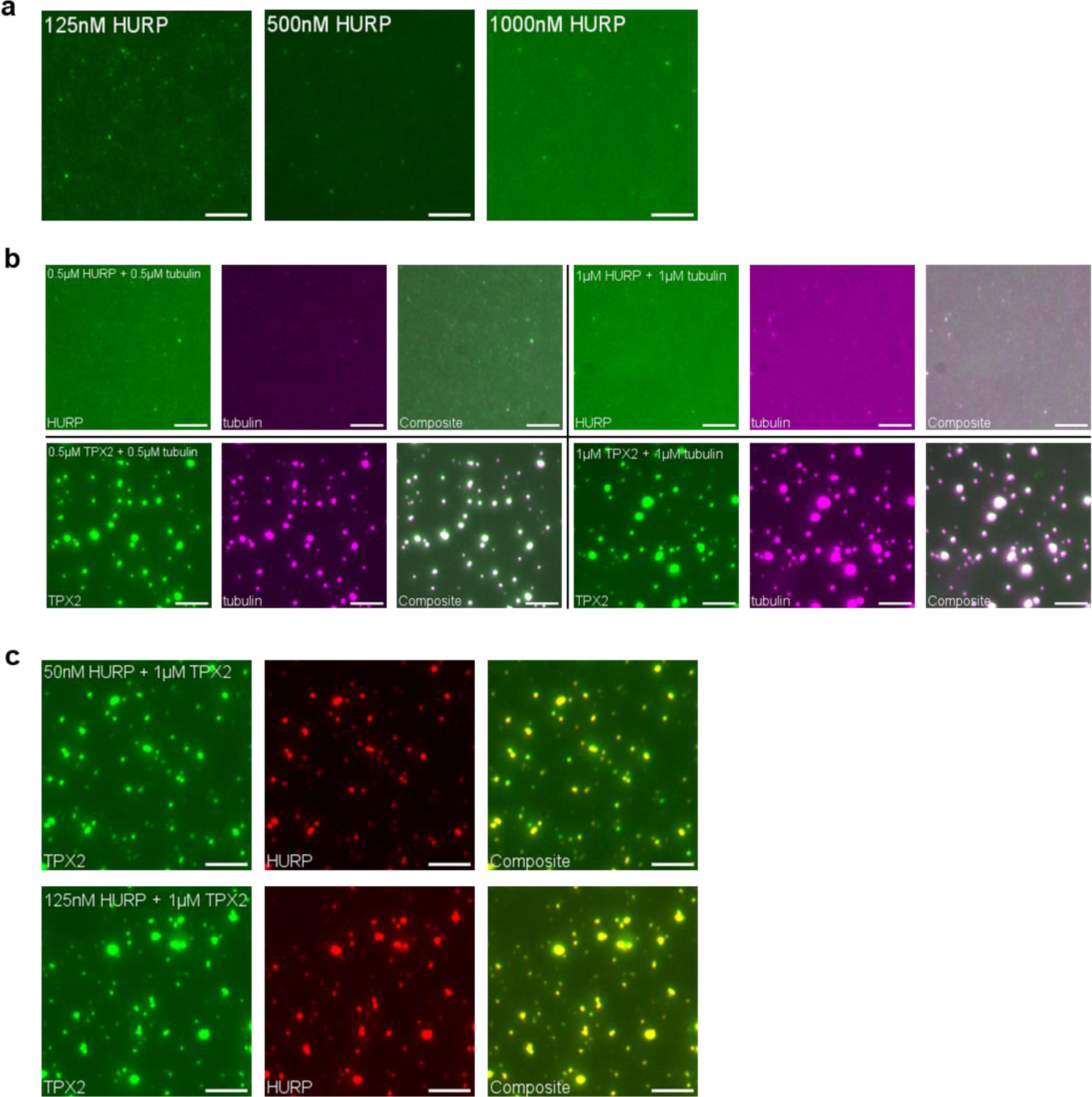
HURP enters TPX2 phase condensates, but HURP alone does not form phase condensates in vitro. **a**Bulk phase assay in BRB80 buffer with increasing concentrations of purified GFP-HURP. **b** Bulk co-condensation assay in BRB80 buffer with equimolar Cy5-tubulin and GFP-HURP at 500 nM and 1 µM (top row). As a positive control, the same experiment was done with GFP-TPX2 (bottom row). Different conditions are separated by black lines. **c** Bulk co-condensation assay in BRB80 buffer with condensing TPX2 (1 µM TPX2) and either 50 nM (top row) or 125 nM mCherry-HURP (bottom row). All images were taken at the surface of the coverslip to better visualize phase droplets. Representative cropped images (15.05 µm x 15.05 µm) are displayed. Scale bars = 5 µm.

**Supplementary Figure 6:**
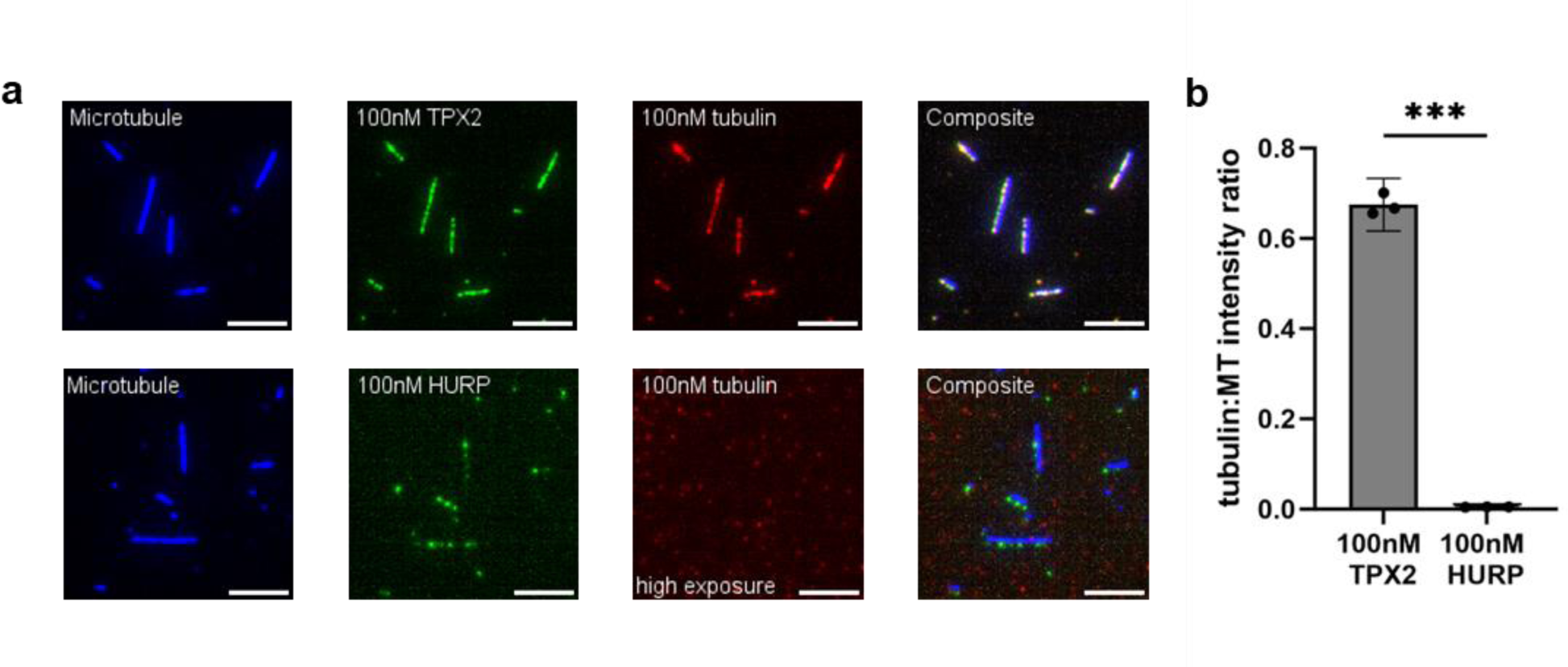
HURP does not directly bind and localize soluble tubulin to pre-existing microtubules. **a**In vitro microtubule (MT) binding reactions using GMPCPP-stabilized and biotinylated MTs bound to coverslips. 100 nM soluble tubulin (Alexa Fluor 568-tubulin) was preincubated with either 100 nM GFP-TPX2 (positive control, n=3) or 100 nM GFP-HURP (n=3) prior to incubating with MTs. Each row of images represents a different reaction. Before imaging, MTs were washed with BRB80 buffer to remove unbound proteins. The tubulin channel for the HURP condition was taken at 25x higher exposure than the TPX2 condition to ensure visualization of any tubulin localization. Representative cropped images (39.77 µm x 39.77 µm) are displayed. MTs (ATTO647N-tubulin) were pseudo-colored blue for display. Representative Scale bar = 5 µm. **b** Bar graph plotting the average tubulin:MT intensity ratio for each MT binding reaction of panel a. For the HURP condition, the tubulin:MT intensity ratio was scaled down by a factor of 25 to account for the exposure difference to the TPX2 condition. Each individual point on the bar graph represents the average intensity ratio for each replicate. For each replicate, 30 or more MTs were measured. Error bars represent the 95% confidence interval. Significance is defined as a p-value < 0.05 by Welch’s two-tailed t-test. Triple asterisks (***) indicates a p-value < 0.001.

**Supplementary Figure 7:**
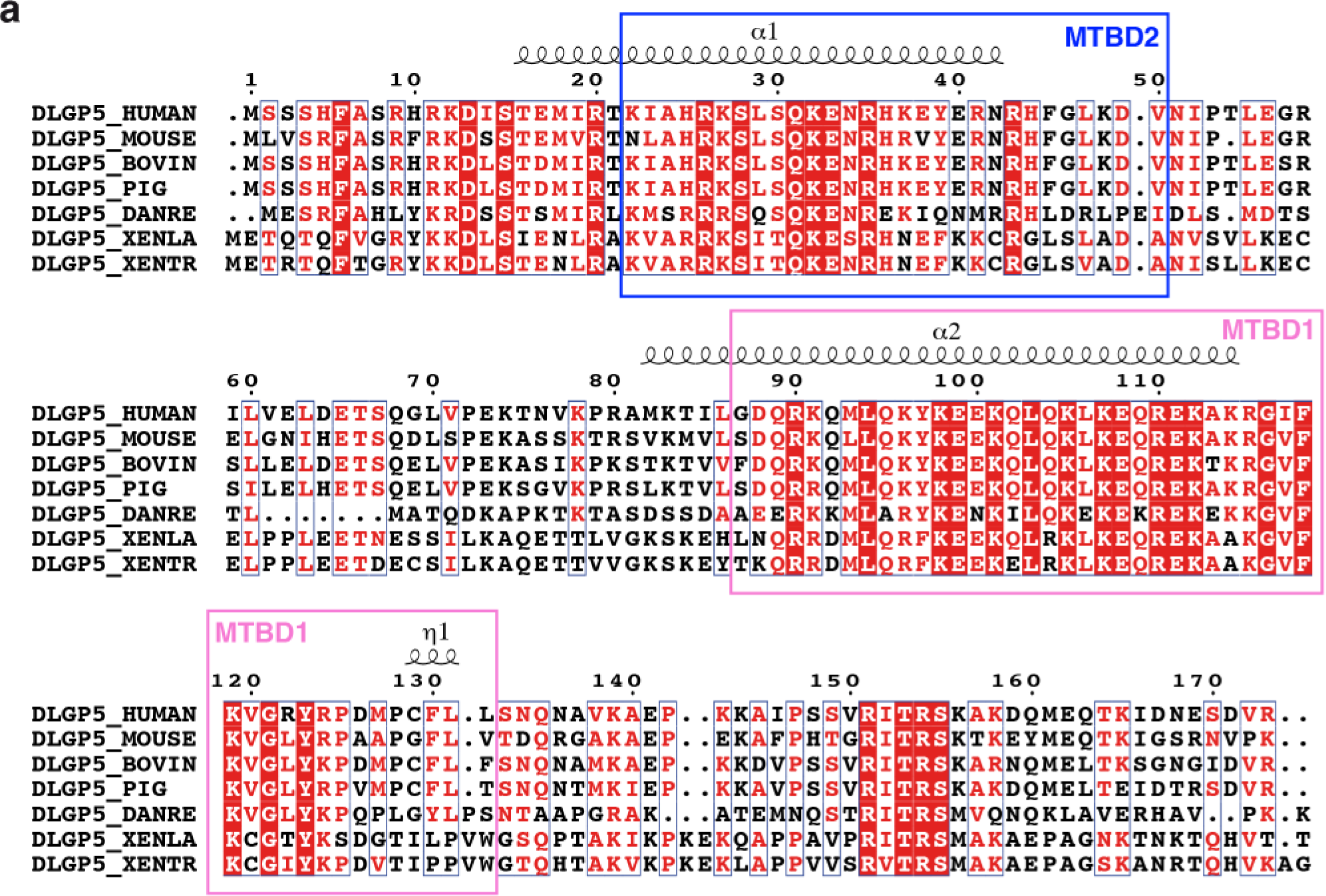
Sequence alignment of HURP from different species. **a**The figure was prepared using ESPript3^68^ based on alignment results from Clustal Omega^69^. Secondary structure elements of HURP from Alphafold2 prediction (Fig. 5) are shown above the sequences.

**Supplementary Figure 8:**
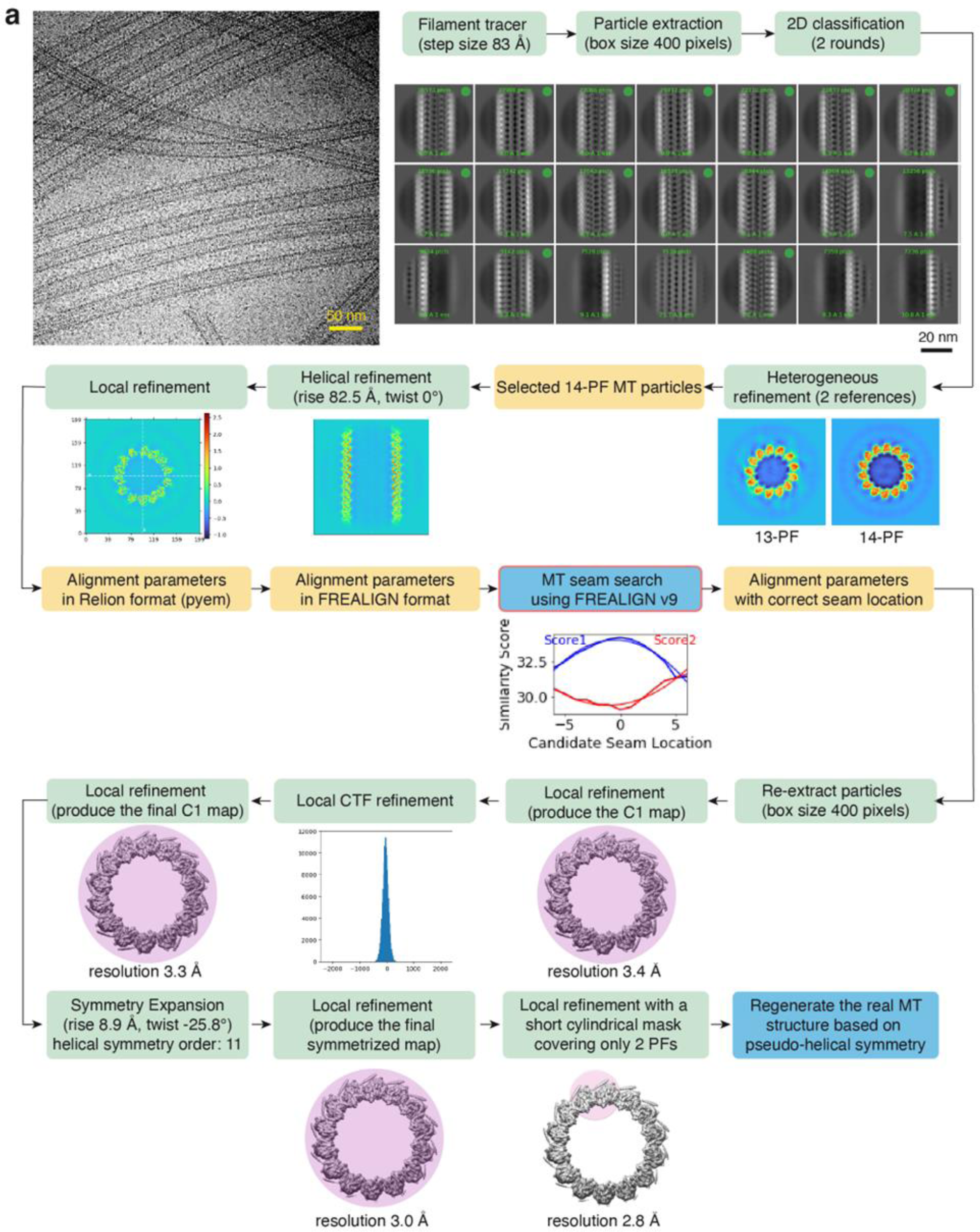
Cryo-EM data processing of HURP decorated microtubule. **a**Functions/operations in CryoSPARC are indicated by green boxes. Cross-sections of the cylindrical masks used for local refinement in CryoSPARC are shown in purple.

**Supplementary Table 1:** Cryo-EM data collection, refinement, and validation statistics.

|  |  |
| --- | --- |
|  | **EMD-XXXXX, PDB: XXXX** |
| **Data collection and processing** |  |
| Microscope | Titan Krios G3 (Thermo Fisher) |
| Detector | K2 Summit (Gatan) |
| Voltage (keV) | 300 |
| Nominal magnification | 81,000× |
| Electron exposure (e<sup>-</sup>/Å<sup>2</sup>) | 39.6 |
| Defocus range set during data acquisition (μm) | -1.0 to -2.5 |
| Pixel size (Å) | 1.39 |
| Symmetry imposed | C1 |
| Initial particle no. (8 nm) | 348,514 |
| Final particle no. (8 nm, symmetry expanded) | 2,936,725 |
| Map resolution (Å) | 2.8 |
| **Model composition** |  |
| Chains | 45 |
| Residues | 8,127 |
| Ligands | 9 GTP / 9 G2P / 18 Mg |
| **Refinement** |  |
| Resolution limit set in refinement (Å) | 2.9 |
| Correlation coefficient (CCmask) | 0.8645 |
| Root-mean-square deviation (bond lengths) (Å) | 0.004 |
| Root-mean-square deviation (bond angles) (Å) | 0.636 |
| **Validation** |  |
| MolProbity Score | 1.71 |
| Clash score | 10.52 |
| Rotamer Outliers (%) | 0.03 |
| Ramachandran (favored) (%) | 97.06 |
| Ramachandran (outliers) (%) | 0.00 |

## Supplementary Materials

**Supplementary Movie 1: HURP is necessary for branching microtubule nucleation.**

Time-lapse movie of Ran-induced branching microtubule nucleation assay in mock-depleted xenopus egg extract, HURP-depleted extract, and HURP-depleted extract with 250 nM purified full-length HURP added back. Reactions imaged over ∼30 minutes. Scale bar = 20 μm. Full 164.92 µm x 139.15 µm field of view shown.

**Supplementary Movie 2: Excess HURP induces microtubule nucleation in *Xenopus* egg extract.**

Time-lapse movie of excess HURP-induced microtubule nucleation in Xenopus egg extract (in the absence of RanGTP) at varying concentrations of HURP. Reactions imaged over ∼30 minutes. Scale bar = 20 μm. Representative 96.63 µm x 96.63 µm cropped field of view shown.

**Supplementary Movie 3: TPX2 and HURP work synergistically to nucleation microtubules in *Xenopus* egg extract.**

Time-lapse movie of microtubule nucleation in Xenopus egg extract induced by addition of excess purified TPX2, HURP, or both together (in the absence of RanGTP). Reactions imaged over ∼30 minutes. Scale bar = 20 μm. Representative 96.63 µm x 96.63 µm cropped field of view shown.

**Supplementary Movie 4: HURP facilitates γ-TuRC nucleation in vitro.**

Time-lapse movie of in vitro γ-TuRC-mediated microtubule nucleation assay in the presence of 50 nM HURP. Reactions imaged over 5 minutes. Scale bar = 5 μm. Full 81.45 μm x 81.45 μm field of view shown.

